# Agent-based modeling demonstrates how target-independent processes supplement killing by antibody-drug conjugates in cancer therapy

**DOI:** 10.64898/2025.12.26.696346

**Authors:** Melissa C. Calopiz, Jennifer J. Linderman, Greg M. Thurber

**Affiliations:** Department of Chemical Engineering, University of Michigan, Ann Arbor, Michigan, United States of America; Department of Biomedical Engineering, University of Michigan, Ann Arbor, Michigan, United States of America; Rogel Cancer Center, University of Michigan, Ann Arbor, Michigan, United States of America

## Abstract

Antibody-drug conjugates (ADCs) have had remarkable clinical success in recent years with multiple new approvals. However, for some ADCs, the response rates don’t closely correlate with clinical target expression. One particular ADC targeting HER2, trastuzumab deruxtecan or T-DXd, is notable due to its success at expression levels ranging from high to low and ultralow. This raises the question of the relative contributions of target-independent mechanisms on ADC efficacy in the clinic, and several such mechanisms have been proposed. However, *in vitro* and preclinical data have different doses and exposures, making it challenging to quantitatively extrapolate preclinical data to the clinic. In this work, we use our computational hybrid agent-based model, *SimADC,* to simulate target-dependent and -independent mechanisms, scaling from mice to humans. We first demonstrate that CD8+ T cells can significantly contribute to tumor regression, especially when the ADC further activates the immune cells. Next, we test target-independent payload-driven mechanisms including: 1) Fc-mediated internalization of ADC by intratumoral macrophages and payload release to neighboring cancer cells, 2) free payload circulating in the blood and re-entering the tumor, and 3) extracellular linker cleavage and payload release due to an abundance of proteases in the tumor. We find that free payload in the blood and extracellular linker cleavage had low and moderate impacts, respectively, while macrophage uptake and payload release resulted in high levels of efficacy. This is due to the macrophages’ ability to sustain free payload in the tumor. Moderate and high HER2 expression were more efficacious than target-independent mechanisms. Overall, our simulations demonstrate that moderate to high HER2 expression, immune activation, or macrophage uptake and payload release are sufficient for T-DXd tumor regression. Additionally, *SimADC* provides a robust framework for modeling both target-dependent and target-independent mechanisms for any ADC, providing the opportunity to engineer more effective therapeutic agents.

**Author Summary:** Cancer is one of the most prevalent diseases in the world, impacting the lives of millions of people every year. Antibody-drug conjugates (ADCs) are a form of targeted therapy that can deliver cytotoxic drugs directly to cancer cells, increasing efficacy. However, ADCs are complex to design and test, as each part of the ADC (targeting antibody, cytotoxic payload, and linker) must be optimally selected for delivery for each target and type of patient. Here, we studied ADCs using a computational model, which allowed us to simulate ADCs in varying cancer environments efficiently and economically. We validated our model using preclinical data to incorporate patient immune responses, target-independent payload release, and systemic payload uptake, allowing us to make accurate predictions in mice and extrapolate to human tumors. We compared multiple mechanisms by which ADCs can kill cancer cells to help identify the most effective methods. Besides high target expression, immune stimulation and target-independent release in the microenvironment can contribute to tumor regression. Investigating these mechanisms enables the design of ADCs and treatment regimens that maximize efficacy across a range of tumor types and target expression.

## Introduction

ADCs have demonstrated significant growth and success in treating solid tumors with six FDA-approved ADCS in the last five years. ADCs bind to specific antigens present on cancer cells, internalize, and release the payload intracellularly for targeted payload delivery to the tumor. Most of these targets are highly expressed in solid tumors relative to low or negligible expression in healthy tissue, and clinical data highlight greater response rates in patients with higher expression levels [1,2]. However, recent data surprisingly show that some ADCs have substantial efficacy in low expression patients. This not only raises questions on their mechanism of action but also provides opportunities to engineer more effective agents.

As an important example, the ADC trastuzumab deruxtecan (FDA-approved Enhertu, or T-DXd) is efficacious across multiple expression levels in breast cancer in the clinic. Recent data, particularly with more sensitive methods than immunohistochemistry (IHC) staining, show stronger responses with higher expression [3]. However, several studies have demonstrated significant benefit at very low or even negative expression levels [4]. Computational analysis shows that the potency and DAR (drug-to-antibody ratio) of T-DXd are optimized for patients moderately overexpressing HER2 (∼100K HER2/cell, similar to IHC2+) [5]. These IHC2+ tumors can be fully saturated by the ADC, resulting in the maximum payload cell killing efficiency [6]. Higher HER2 expression (∼1 million HER2/cell or IHC3+) result in poor tissue penetration, but bystander effects (free payload diffusing out of targeted cells into neighboring untargeted cells) partially compensate for this effect, resulting in strong efficacy. However, at low expression levels, trastuzumab-based ADCs can quickly saturate tumors, but not enough payload enters the cell even with cell saturation, as seen with FDA-approved Kadcyla with lower HER2 expression [2,7]. This can result in reduced efficacy. However, some patients show significant responses to T-DXd even with low and/or undetectable HER2 expression with the methods used. It is currently unknown why T-DXd is efficacious at these lower expression levels in breast cancer.

Several mechanisms have been proposed to drive the efficacy of T-DXd at low expression levels (**Fig 1**). One hypothesis is that the immune system contributes additional cell killing, specifically mediated through CD8+ T cells. In preclinical models, ADC efficacy is greater in immunocompetent mice versus immunodeficient mice, supporting this possibility. Immune cell activation markers are increased in these models, but it is unclear if the improved efficacy is due to the presence of T cells independent of the ADC (additivity) or higher activity from the activation of T cells by the ADC (synergy) [8,9]. A second hypothesis posits payload efficacy driven by myeloid cells. Myeloid cells can play an important role in payload release through Fc binding, where cells such as macrophages bind to, internalize, and degrade ADCs through their Fcγ receptors, releasing free payload into the tumor to be taken up by cancer cells via bystander effects (hereafter referred to as *mac release*). Literature shows an improvement in ADC efficacy in xenografts with high counts of macrophages, specifically through Fc binding, even for ADCs without target binding in the tumor [10]. Third, cleavage of ADCs due to high levels of extracellular proteases can also release payload independent of the target expression (hereafter referred to as *tumor microenvironment (TME) release*) [11]. This mechanism has also been leveraged for binding slow or non-internalizing targets [12]. The fourth hypothesis includes the systemic release of free payload from the ADC, either by deconjugation in circulation or ADC metabolism in healthy tissue and release of the payload out of the tissue into the circulation (hereafter referred to as *systemic release*). The low but persistent plasma concentrations of free payload may contribute to efficacy through sustained exposure to the small molecule payloads [13].

**Fig 1.**
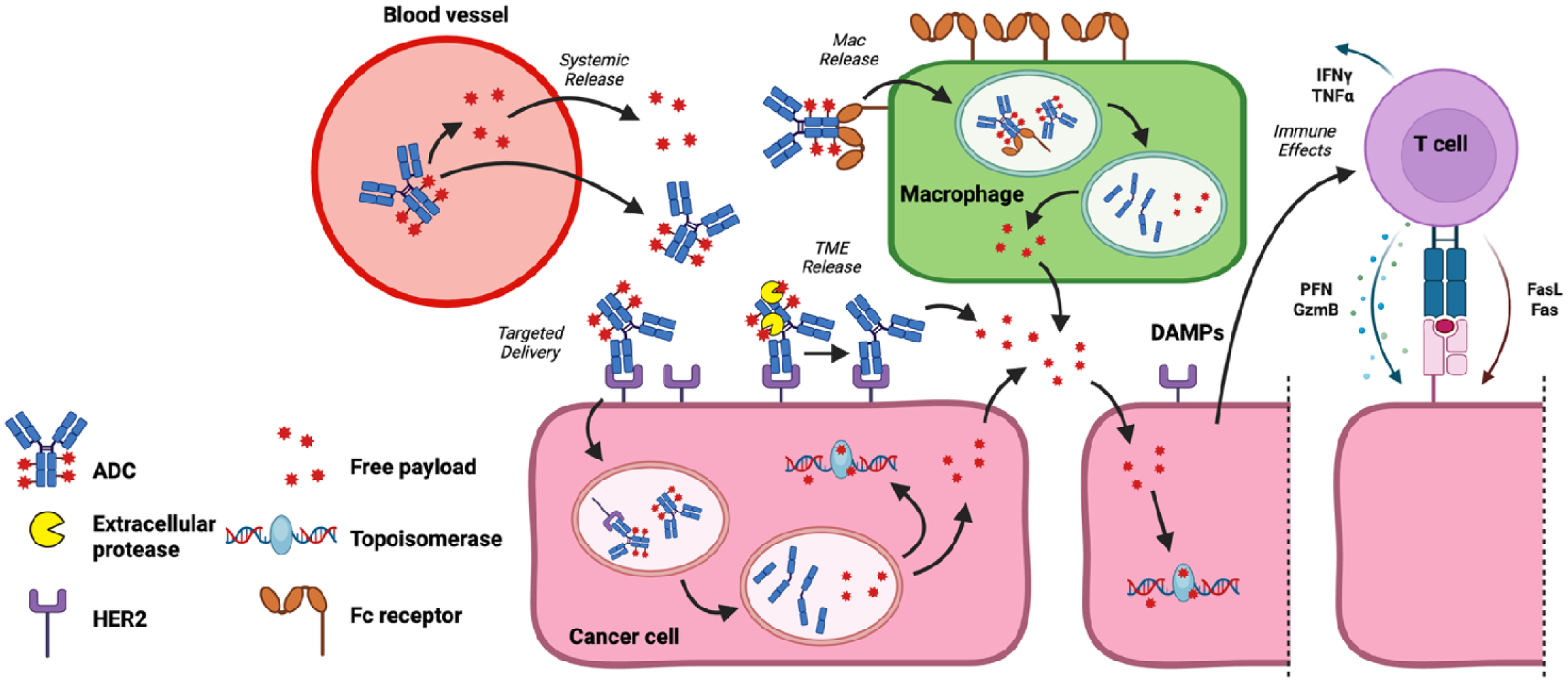
Overview of ADC mechanisms of action. Intact ADCs extravasate from the blood into tumor tissue to enter the tumor microenvironment (TME). The intact ADC typically binds to the cancer cell, internalizes, and releases the payload inside the cell. The payload can directly kill the cell or diffuse into nearby cells for bystander effects. These are generally considered *Targeted Delivery*. Four proposed target-independent mechanisms of efficacy are also shown: (1) Free payload from systemic deconjugation or metabolism in healthy tissue and intravasation can also enter the tumor as free payload (*Systemic Release*). (2) Fcγ receptor mediated uptake of ADCs, often driven by FcγR1, releases payload inside myeloid cells like macrophages (*Mac Release*), and the payload can kill nearby cancer cells (bystander killing). (3) Extracellular proteases can cleave the linker to release payload outside of cancer cells, relying on the bystander effect to enable the payload to enter cells (*TME Release*). (4) An indirect mechanism of efficacy is immune stimulation, which could be driven by mechanisms such as immunogenic cell death, in which cancer cells killed by targeted delivery release damage-associated molecular patterns (DAMPs) to activate T cells for further cancer cell killing. Activated T cells can kill cancer cells by releasing perforin (PFN) and granzyme B (GzmB), Fas/FasL binding, and interferon gamma (IFNγ) and tumor necrosis factor alpha (TNFα) cytokine release to induce apoptosis (*Immune Effects*). The activation of T cells can enhance/drive tumor efficacy depending on the immune microenvironment. Image generated in BioRender.

One of the challenges with distinguishing the quantitative contribution of these ADC mechanisms is the difference between *in vivo* mouse experiments and clinical results. First, the tolerability of ADC payloads can vary significantly between mouse cells and human cells; mouse models generally tolerate higher doses than humans. This can result in artificially large therapeutic windows in preclinical models [14]. Additionally, the relationship between ADCs and the immune system is relatively understudied. Immunodeficient models like xenograft mouse models use human cancer cells that mimic payload sensitivity but lack full immune effects. In contrast, immunocompetent models like syngeneic models capture innate and adaptive immune responses, but they typically use mouse cancer cells that may be less sensitive to the payload, lacking the capacity to capture payload potency in human cancer cells. Therefore, the relative impact of these mechanisms of action is dependent on the animal model chosen, leading to intrinsic challenges in interpreting and translating the data. Differences in pharmacokinetics can also contribute. Mice clear free payload from the plasma much faster than humans, reducing the effects of free payload in the plasma in mice versus humans.

Given these challenges, it is difficult to compare the relative clinical magnitude of various mechanisms experimentally. We thus turned to computational modeling. In this work, we expand our hybrid agent-based model (ABM), *SimADC*, to capture these target-independent mechanisms of action, building on preclinical and clinical data sets to capture immune cell efficacy, mac release, TME release, and systemic release. We then simulate the impact on tumor growth inhibition for individual and combinations of mechanisms with parameters scaled to human patients to provide insight on the relative contribution of these target-independent effects for T-DXd. These results can be used to interpret clinical data, formulate clinical hypotheses, and guide the design of more effective agents.

## Results

*SimADC* is a hybrid ABM that describes the delivery and action of ADCs within a solid tumor. The model is a hybrid between deterministic ordinary and partial differential equations to model ADC and free payload distribution within the TME and a probabilistic representation of thousands of cellular ‘agents’ that can grow and respond to this microenvironment (**Fig 2**). These individual cells can divide, die, or in the case of immune cells, kill cancer cells, according to probabilistic rules. Thus, simulations are capable of describing the time and spatially varying pharmacokinetics and pharmacodynamics (PKPD) of ADCs in solid tumors.

**Fig 2.**
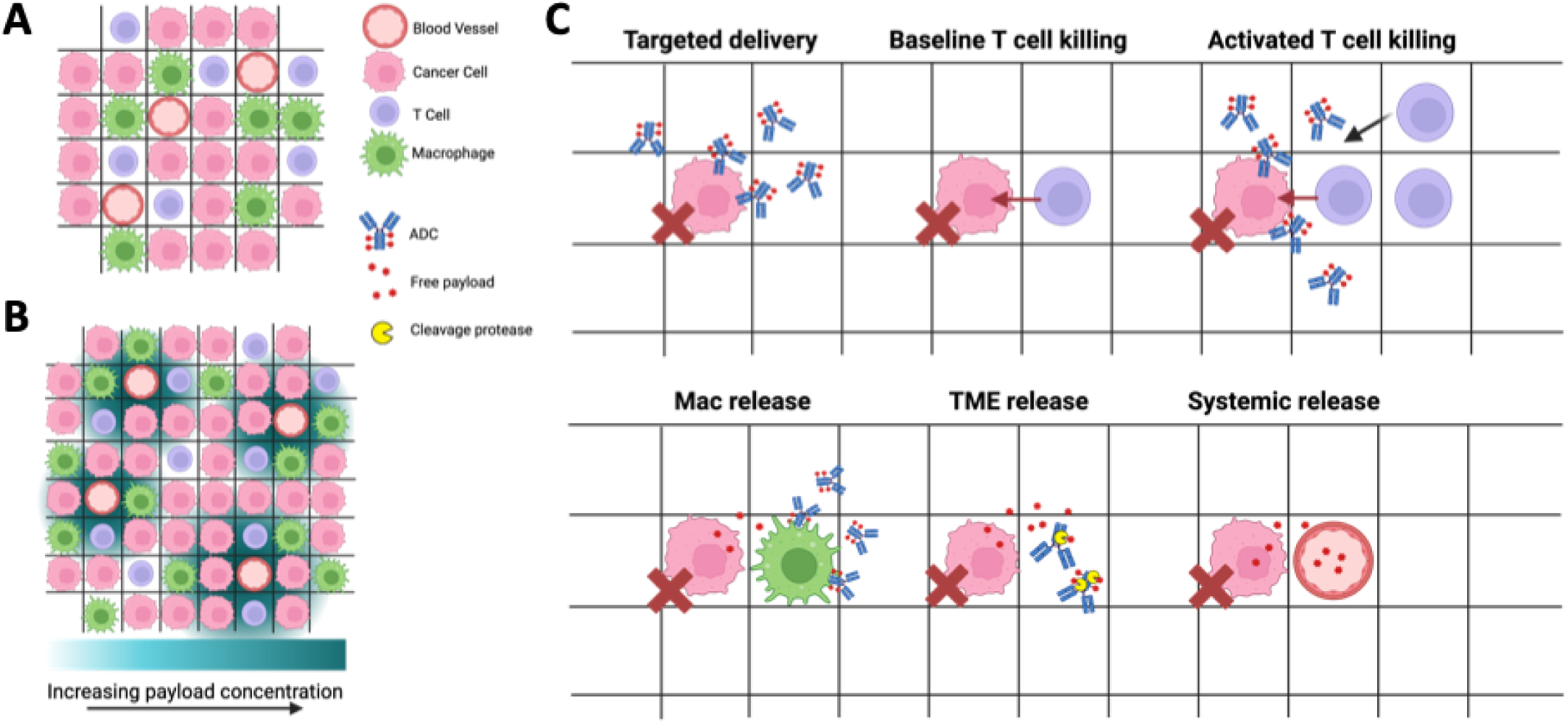
Hybrid Agent-Based Model (*SimADC)* Combines Drug Gradients with Cellular Agents. (A) A grid of thousands of cells incorporates blood vessels (source and sink of ADCs and payloads from the plasma), cancer cells that divide or die with probabilities based on local conditions, T cells for immune cell killing, and macrophages for Fc-mediated payload release. (B) The grid is overlayed with drug gradients (including bound and unbound ADC and free payload in the intracellular and extracellular space) that are used to capture bystander killing and determine the probability of cell death. (C) The updated model incorporates targeted delivery (inclusive of bystander effects) as in our previous models but also adds T cell killing (baseline and activated), macrophage-mediated payload release (mac release), extracellular ADC linker cleavage (TME release), and uptake of free payload from the plasma (systemic release). Image generated in BioRender.

In this work, we expanded our model by adding (1) baseline and activated T cell killing, (2) mac release (**Fig S1**), (3) TME release, and (4) systemic release. Each mechanism was added into *SimADC* and calibrated to literature data to capture the effects in mice. Then, the mechanisms were scaled to clinically relevant doses and concentrations in humans for quantitative predictions of clinical effects.

All model equations and parameters are included in Methods.

### *SimADC* captures efficacy from targeted delivery and baseline T cell killing independently

ADCs are more effective in immunocompetent mouse models than immunodeficient mouse models [8,9], so a data set capturing both payload and immune effects was needed to train our model. Rios-Doria et al. [8] published efficacy data for two ADCs, EphA2-Tubulysin and EphA2-PBD, in a CT26 mouse model in both nude and syngeneic mice, providing data to train the ABM model. First, the cell doubling time was fit to match the intrinsic growth rate of the untreated tumor. Then, the pharmacodynamic parameters K_m_ and P_max_ (see Methods) for each payload were fit to capture the distribution and efficacy of each payload in CT26 tumors grown in nude mice. The dose response curves closely match the experimental efficacy data (**Fig 3A**). While many payloads have low efficacy in mouse cells like CT26, the Tubulysin and PBD agents show moderate to high potency in these cells.

**Fig 3.**
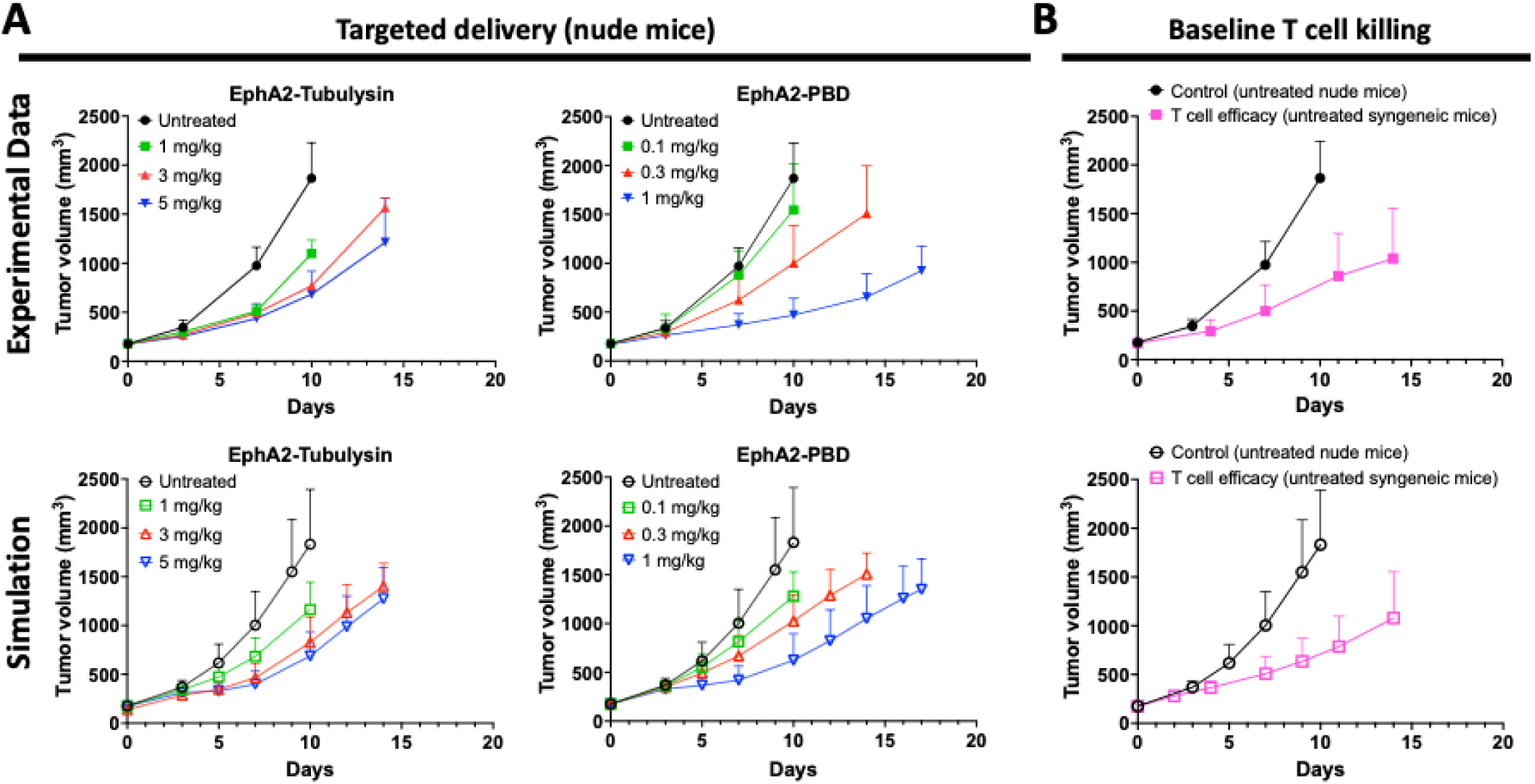
Independent calibration of payload potency and baseline T cell activity. Experimental data from Rios Doria et al. [8] (top row) were used to train the simulations (bottom) on payload efficacy and T cell effects. (A) The cell doubling time was fit to match the untreated animal model tumor growth, and pharmacodynamic parameters were fit to efficacy data using an anti-EphA2 antibody with either a Tubulysin or PBD payload in a nude mouse model that lacks T cell effects. *SimADC* was able to capture the magnitude and dose response of these effects. (B) For T cell killing, the growth rate in a nude mouse host versus immunocompetent syngeneic host was compared. The baseline (non-activated) T cell killing rate was fit to account for the slower growth rate when intratumoral T cells are present. Error bars represent standard deviation; each simulation point is the result of 300 simulation runs (100 simulations run in triplicate).

To incorporate interactions between ADC treatment and the immune system, we added CD8+ T cells to the TME. CD8+ T cells (hereafter referred to just as T cells) are one of the main effector cells for immune anti-tumor responses and can directly contribute to tumor regression. Data from Rios-Doria et al. [8] shows that T cell depletion reverses the slower tumor growth of CT26 cells in an immunocompetent mouse model to the faster growth of an immunodeficient model, indicating this is a critical cell type to include in the ABM (**Fig S2**). To incorporate the potential ability for T cells to be activated by treatment, we included two states of the T cell agents, active and inactive, with distinct probabilities of cancer cell killing. The number of and active fraction of T cells in untreated tumors (10%) were estimated based on the reported T cell density and reported activation markers, and we calibrated the T cell killing probabilities to match the reduced growth rate of CT26 cells in an immunocompetent host (syngeneic model). By incorporating a probability of the T cell to kill adjacent cancer cells, the different growth rates in the two hosts was captured (**Fig 3B**). The calibrated model was next used to understand the relative roles of targeted delivery vs. T cell killing, and if targeted delivery can influence T cell killing through activation.

### ADC treatment in immunocompetent mice shows a synergistic response from immune activation

With the drug pharmacodynamic parameters and T cell probabilities independently calibrated, we can simulate the combined impact of ADCs and T cells on tumor regression. When combining independent mechanisms of action (such as payload and immune cell killing), a key question is whether the mechanisms are additive, synergistic, or antagonistic. The increased efficacy seen in immunocompetent hosts could reflect simultaneous targeted ADC delivery and baseline T cell killing (additive), the ADCs could kill a large fraction of T cells (antagonistic), or the ADCs could activate the immune cells (synergistic). We first simulated the efficacy of ADC treatment in an immunocompetent host by simply adding the T cells to the ABM and treating with the ADC – an additive model with no activation – to compare with the experimentally measured efficacy.

The simulated efficacy from the additive payload effects and baseline T cell effects showed a consistent under-prediction of efficacy relative to the experimental data (**Fig S3**). These cell-killing mechanisms are additive at the mechanistic level, but they would include overlapping killing, such as T cells killing a perivascular cell that already internalized a lethal dose of payload, similar to the Bliss model of independence [15]. The additive mechanism resulted in large, biased residuals (**Fig S4**). Consistent with the elevated activation markers seen in these mice, these results indicated that efficacy was greater than additive. We then updated the percentage of active T cells based on values reported from Rios-Doria et al. [8] for one dose of each ADC (18% for 5 mg/kg of EphA2-Tubulysin and 22% for 0.3 mg/kg of EphA2-PBD) to represent a synergistic mechanism, in which the percentage of active T cells increases with ADC dosing. When we simulated each dose of ADC, the synergistic mechanism resulted in tumor volumes closer to the immunocompetent experimental efficacy curves (**Fig 4**). When comparing the residuals of the additive and synergistic mechanisms, we find smaller residuals and relative errors for the synergistic mechanisms (**Fig S4**). Based on these data, the model simulates the efficacy of ADCs with T cell activation by incorporating the increase in active T cells.

**Fig 4.**
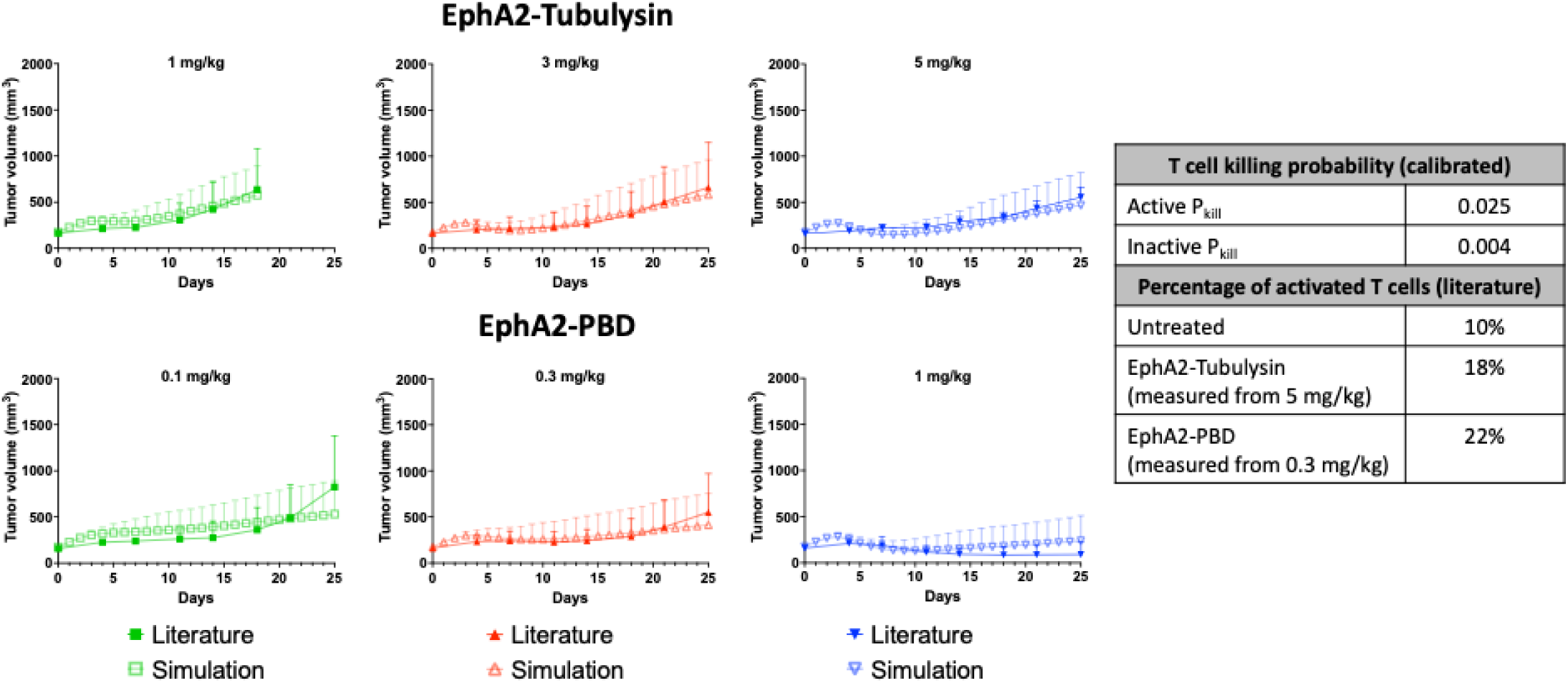
Experimental and simulated efficacy of ADCs in immunocompetent mice. To capture the activation of immune cells following ADC treatment, the percentage of active T cells in the model was increased from 10% (untreated) to 18% (EphA2-Tubulysin) or 22% (EphA2-PBD) based on values reported in Rios-Doria et al. [8]. While only measured for one dose of each ADC (5 mg/kg for EphA2-Tubulysin and 0.3 mg/kg for EphA2-PBD), these values were kept constant across doses. Error bars represent standard deviation; each simulation point is the result of 300 simulation runs (100 simulations run in triplicate).

### T-DXd achieves efficacy in an immunocompetent mouse model largely by immune activation

We next extended these results from the DNA-damaging (PBD) and microtubule inhibitor (Tubulysin) ADCs to topoisomerase inhibitors. Notably, the topoisomerase payload has significantly lower potency in mouse cells than human cells. This can be seen with the lack of efficacy from 10 mg/kg of T-DXd in CT26-hHER2 cells grown in nude mice (**Fig 5**). However, the payload is still able to activate immune cells [9,16].

**Fig 5.**
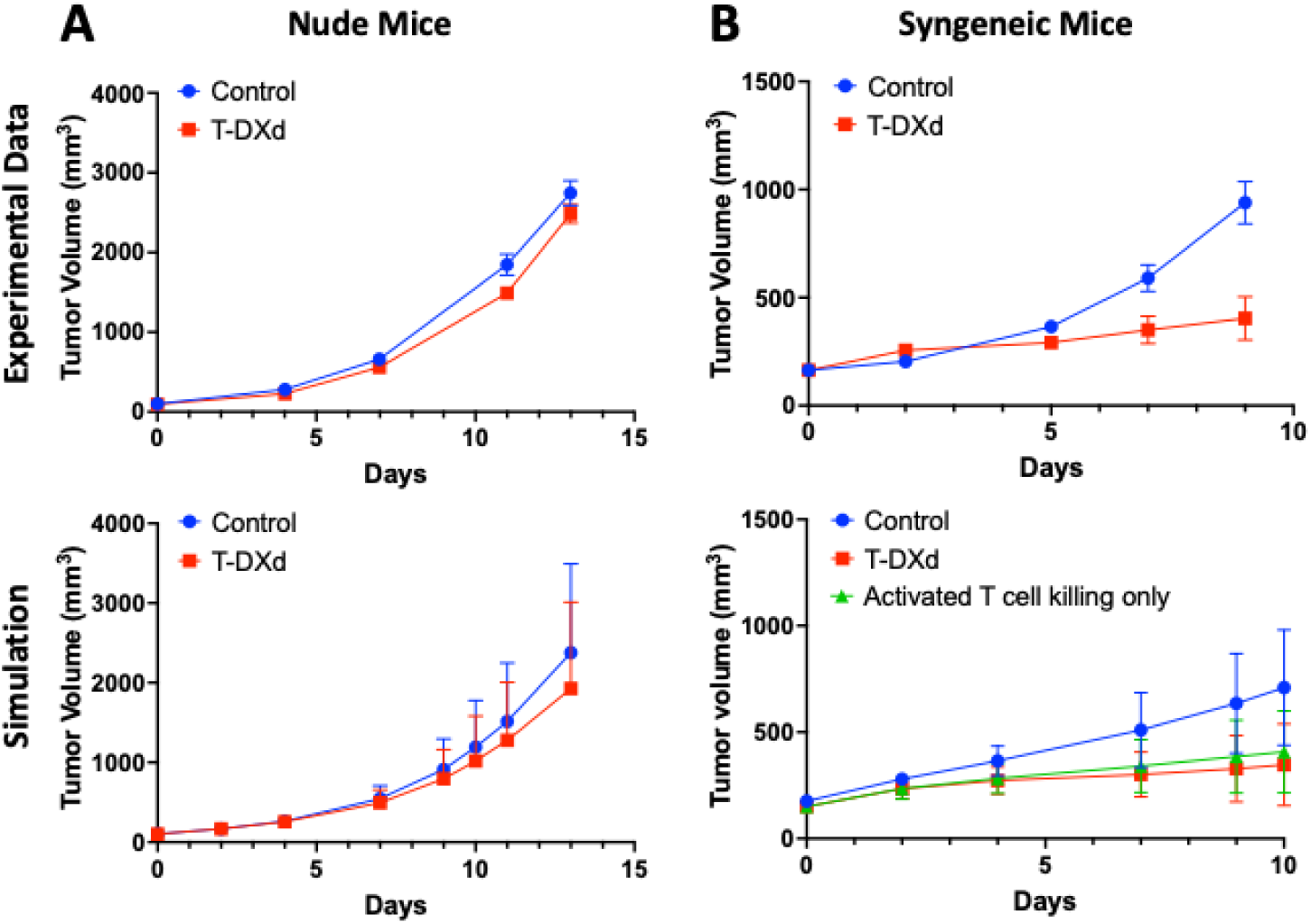
Treating nude vs. syngeneic mice with T-DXd. (A) Using the CT26 nude mice growth rates previously calibrated from Rios-Doria et al. [8], we simulated T-DXd with a 20-fold decrease in potency (to account for the differences in sensitivity between human NCI-N87 cancer cells and mouse CT26 cancer cells) and find agreement with the experimental data showing little efficacy in mouse cancer cells grown in the nude mouse model. (B) Using the growth rates from the syngeneic mouse model, we simulated T-DXd treatment and find a similar efficacy to the experimental data. We also simulated the impact of activated T cell killing alone (no payload efficacy) and see similar tumor volumes, indicating that efficacy is mainly driven by T cell activation in this model. Error bars represent standard deviation; each simulation point is the result of 300 simulation runs (100 simulations run in triplicate).

Utilizing the same T cell density and activation rate from the earlier calibration (**Fig 4**), we show that the efficacy of T-DXd in an immunocompetent host is largely driven by the activation of immune cells. The same simulation with no immune cell killing shows the negligible efficacy seen experimentally with this same cell line grown in the nude mouse model that captures primarily payload effects with minimal immune cell killing (**Fig 5**). These results highlight a challenge in testing the efficacy of ADCs, since efficacy in immunodeficient mouse models is driven by payload killing of cancer cells, potentially underestimating immune effects, while efficacy in immunocompetent mouse models is heavily dependent on immune cell killing of cancer cells while potentially underestimating payload killing due to the lower payload potency.

### Fc-mediated internalization of ADCs captures untargeted ADC efficacy

We next incorporated mac release (**Fig 2C**) to determine its contribution to tumor regression in mice. Aside from the direct impact that immune cells can have on cancer cells, such as the T cell killing incorporated above, cells like macrophages can also interact with the ADC to facilitate payload release. While fluid phase uptake from mechanisms like phagocytosis can contribute, Fcγ-receptor mediated internalization and release of bystander payloads is quantitatively more significant [10]. We included macrophage agents on the grid with the ability to bind, internalize, and degrade ADCs based on measured Fcγ-receptor expression levels and internalization rates [17]. The macrophage rates of ADC binding and internalization are distinct from those of cancer cells and based on Fcγ receptor expression and affinity. The intracellular free payload can then diffuse into adjacent cells and exert bystander killing, as with free payload originating in cancer cells.

For model validation, we used data from Li et al. [10] who reported *in vivo* efficacy and intratumoral drug concentrations from *in vivo* tumor slices for two ADCs, αCD30-MMAE (targeting) and hIgG-MMAE (non-targeting), both at 2 mg/kg. These results are in L428 tumors, which are reported to have high levels of macrophages. By using macrophage pharmacokinetic parameters for IgG from the literature [17,18], we were able to validate our pharmacokinetic parameter set by calculating the intratumoral concentration of our simulated tumors and comparing with the intratumoral concentration data from Li et al. **Fig 6A** shows how the tumor concentration of payload can be similar for both targeted and untargeted antibodies in tumors with large myeloid cell populations.

**Fig 6.**
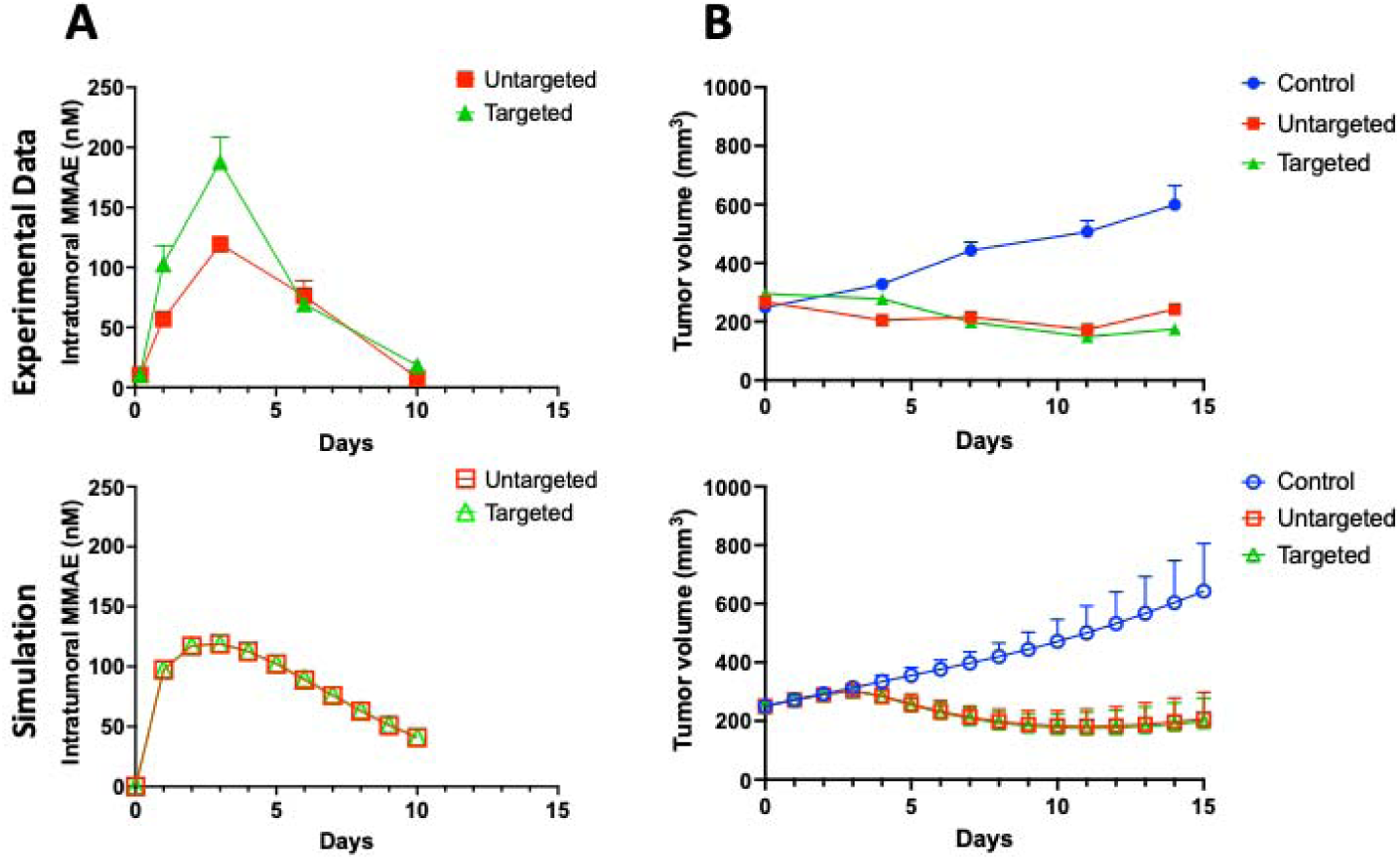
Fc receptor-mediated internalization of ADCs can drive high efficacy in macrophage-dense tumors. Our macrophage model pharmacokinetic and pharmacodynamic parameters were calibrated to literature experimental data from Li et al. [10], who measured (A) intratumoral free payload concentrations and (B) efficacy in tumors with high macrophage infiltration (∼50% macrophages in the tumor) after dosing a targeted (CD30 receptor binding) and a non-targeted (IgG, mainly binding to Fc receptors on macrophages) ADC with an MMAE payload. Our simulations show similar concentrations of intratumoral payload, also resulting in similar efficacy with both types of ADCs after model calibration. Error bars represent standard deviation; each simulation point is the result of 300 simulation runs (100 simulations run in triplicate).

For *in vivo* calibration of efficacy, the tumor growth rates were fit to the control (untreated) tumor and the PD cell killing parameter was calibrated for both ADCs (**Fig 6**). To simulate the targeted ADC, we used a receptor expression for the cancer cells of 108,000 CD30/cell from the literature [19] thereby including CD30 expression with our macrophage parameter set. To simulate the non-targeting ADC, we set the receptor expression for cancer cells to zero to represent the impact of only macrophage uptake on tumor regression. Our fits indicate that the model was able to capture the impact of macrophages on tumor regression.

Given the large concentration of macrophages in these tumors (50% of the tumor) and high level of FcγR expression (∼400,000 receptors/cell), we also simulated a reduced number of macrophages. **Fig S5** shows that the simulated efficacy in tumors with 5% myeloid cells but 775,000 Fcγ receptors per cell is similar to tumors with 10% myeloid cells with half that number of Fc receptors per cell. This indicates the total number of receptors (number of macrophages and receptors/macrophage) correlates with the response. Even with a smaller percentage of macrophages expressing 200,000 – 400,000 Fc receptors/cell, we are still able to achieve a reduction in tumor volume with 10-20% tumor macrophages in these simulations.

### TME release yields efficacy in antigen-negative tumors

TME release (**Fig 2C**) could also contribute to efficacy for T-DXd in some antigen-negative models [11]. To evaluate the magnitude of efficacy from TME release, we included a parameter (k_Cleave_) and additional term in the differential equations (see Methods). This value represents the rate constant at which enzymes could release payload from unbound and bound extracellular ADC in the tumor, converting it to unbound and bound antibody plus extracellular free payload. Because *in situ* cleavage rates in the TME are difficult to measure, this parameter was calibrated to data from Tsao et al. [11], who reported efficacy data for T-DXd in two experiments using MDA-MB-468, a HER2-negative mouse cell line that expresses extracellular proteases like cathepsin L (CTSL). It was not possible to independently evaluate contributions from macrophage uptake, so the efficacy was assumed to be completely from extracellular proteolysis, giving us a possible overestimate of the TME release contribution. In this xenograft model in a SCID mouse host, they observed significant efficacy with T-DXd following 3 weekly doses of 10 mg/kg. Efficacy was also seen in this xenograft model in NOD-SCID mice by Vasalou et al. [20] but not by Ogitani et al. [21] in a nude mouse model, indicating the TME and/or mouse strain can impact the target-independent response. For both efficacy curves, a value of 4×10^-5^/s for k_Cleave_ resulted in the best fit, equating to a cleavage half-life of about 5 hours (**Fig 7**). With this fitted parameter, the efficacy from TME release can be simulated for T-DXd at different dosing regimens and clearance rates (e.g. mouse versus human). The simulated tumor growth response matches the experimental results when treating small and moderated-sized tumors.

**Fig 7.**
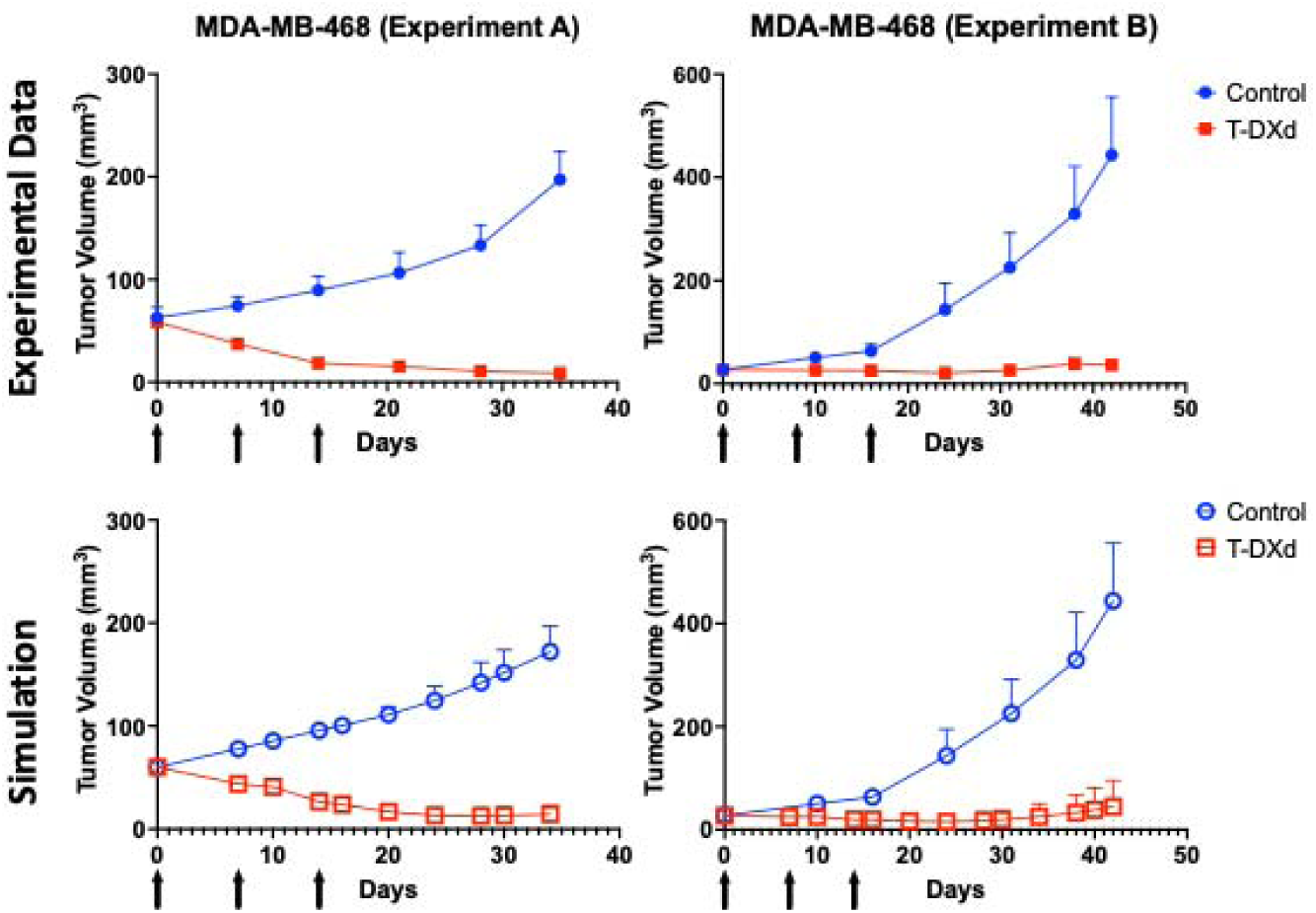
TME release of payload can drive efficacy in tumor xenografts. Experimental data from Tsao et al. [11] (top) shows the growth of untreated HER2-negative MDA-MB-468 xenografts injected in the mammary fat pad of SCID mice (control) or treated with three injections of 10 mg/kg T-DXd on days 0, 7, and 14. Experimental data (Experiment A) were collected from a mouse with HER2-positive and HER2-negative dual-implanted tumors and data (Experiment B) were collected from a mouse with only one HER2-negative tumor. In simulations, a cleavage rate constant (k_Cleave_) of 4×10^-5^/s was able to capture the efficacy in the HER2-negative tumor. Error bars represent standard deviation; each simulation point is the result of 300 simulation runs (100 simulations run in triplicate).

### *SimADC* can predict the efficacy of small molecule chemotherapy with multiple dosing regimens

Clinical data demonstrate that the concentration of free payload in the blood resulting from ADC administration can be higher than the *in vitro* IC50, making it plausible that this also contributes to efficacy. The free payload originates from the detachment of the ADC in circulation or intact ADC uptake within healthy tissue that releases the payload, enabling it to diffuse back into the blood, where it can enter the tumor. Importantly, the clearance of free payload is faster in mice than in humans, so the rapid clearance in mice results in much lower plasma payload concentrations. Therefore, this effect would not show up in preclinical models of ADC efficacy. The slow but continuous release of small molecule chemotherapy is in line with other work aimed at maintaining low but consistent levels of chemotherapy drugs in the plasma. Some of these similar studies include continuous intravenous infusion (CIVI) of small molecule drugs [22]. To capture the magnitude of this effect (systemic release, **Fig 2C**), we first needed to demonstrate that *SimADC* can be used to simulate efficacy from small molecule concentrations in the blood, since thus far it has only been validated for the delivery of biologics.

For small molecule drug delivery to tumors, notably, these agents are blood-flow limited rather than permeability-limited [14,23,24]. To account for blood flow-limited delivery, we modified the effective blood vessel permeability rate based on vessel depletion data seen in the clinic [25]. While small molecules can extravasate quickly by paracellular and transcellular diffusion, we reduced the effective permeability 3-fold so the net tumor uptake rate was equal to the blood-flow delivery rate of small molecules into tumors. This allowed us to use the 2-dimensional approximation of the tumor microenvironment rather than a 3D simulation accounting for axial decreases in tumor blood concentration. Using plasma pharmacokinetic data from intravenous administration of free DXd [26], we matched the systemic clearance rates using a non-linear regression model for two phase decay and determined the equivalent free payload dose to use in the model (**Fig 8A**). To validate our new PK and existing PD parameter sets, we simulated several multi-day dosing regimens of free exatecan as published in Kumazawa et al. [27] (**Fig 8B**). We adjusted our PD parameter set to account for the change in potency between exatecan and DXd (exatecan is almost twice as potent as DXd), scaled the experimental dose to our equivalent free payload dose matching the plasma pharmacokinetics, and simulated the delivery of free payload. The results show that *SimADC* is able to predict the efficacy from systemic small molecule chemotherapy to each dosing regimen. More importantly, the model can simulate efficacy from free payload in the blood to predict the contribution to ADC efficacy by incorporating the systemic free payload concentrations.

**Fig 8.**
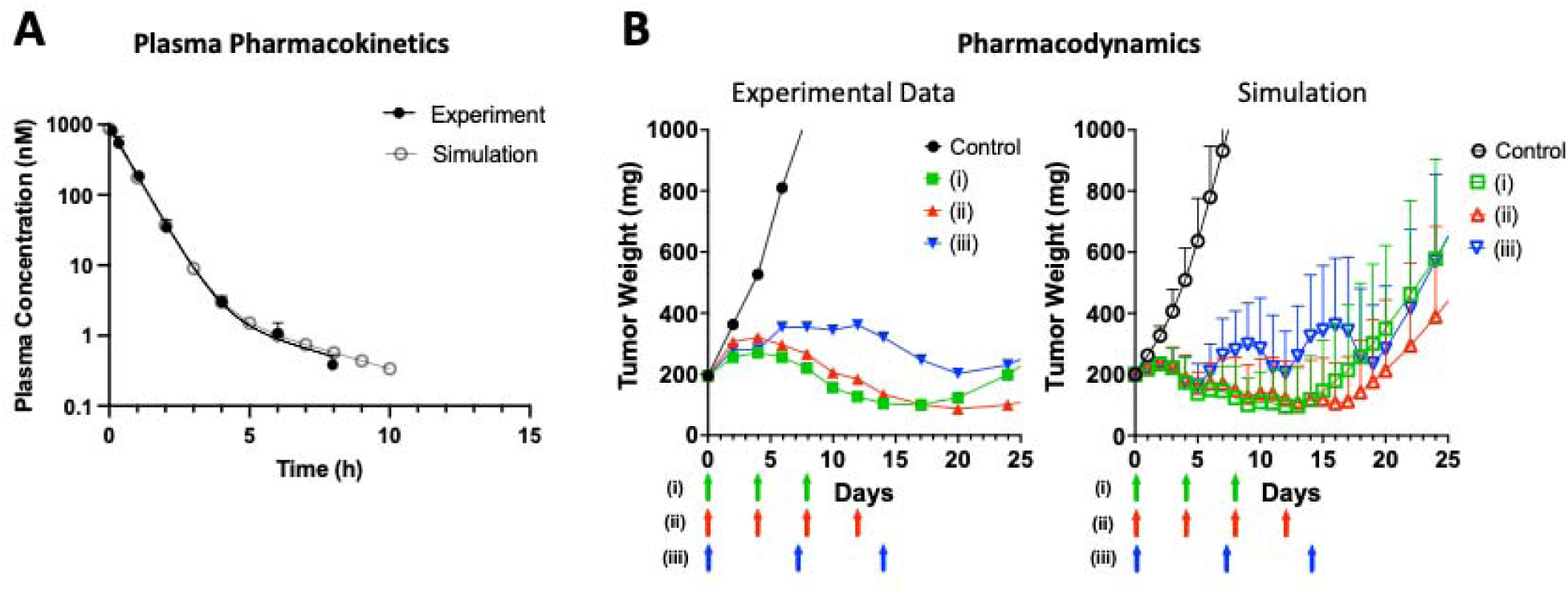
*SimADC* predicts responses to systemic DXd in mouse xenografts. (A) The plasma clearance parameters for free DXd were fit to experimental data from Okamoto et al. [26] following intravenous delivery. We simulated a dose of 0.04 mg/kg (vs. 1 mg/kg used in the experiment) to match the initial concentration. (B) Our previously-calibrated DXd parameter set was validated by comparison to experimental data from Kumazawa et al. [27] (left) after calibrating our cell doubling time parameters to the control curve measured from human SC-6 xenografts in nude mice. We doubled our maximum probability of cell killing to adjust for the change in potency (0.045 for DXd vs. 0.09 for exatecan) and dosed a total of 2 mg/kg (vs. a total of 50 mg/kg used in the experiment) based on our equivalent model dose from (A). We simulated the following dosing regimens as outlined in Kumazawa et al.: (i) q4d x 3, (ii) q4d x 4, and (iii) q7d x 3. Error bars represent standard deviation; each simulation point is the result of 300 simulation runs (100 simulations run in triplicate).

### *SimADC* predicts a minimum of 30,000 HER2/cell in patients for efficacy from targeted delivery alone

To understand the clinical relevance of these incorporated mechanisms, we next turned to understanding human data by scaling up our pharmacokinetic parameters to match human clearance rates. We simulated varying HER2 levels with both mouse and human pharmacokinetics (see Methods) to identify the homogeneous expression level needed for payload efficacy alone. T-DXd is approved for HER2-positive, HER2-low, and HER2-ultralow metastatic breast cancer. While clinical data show greater responses in patients with higher expression, there is great interest in quantifying the mechanisms responsible for efficacy in HER2 low or negative expression (with limitations in expression measurement discussed later). Previous work has demonstrated that direct delivery by cancer cell binding and internalization is more efficient than other mechanisms of payload delivery (such as bystander effects) [28,29]. Therefore, we first simulated the minimum HER2 expression needed to drive target-dependent efficacy without other contributing mechanisms. For a threshold, we used a previously published correlation between ADC clinical approvals and the ability to shrink a tumor over the clinical dosing window (e.g., 3 weeks for T-DXd) with a single administration at the clinical mg/kg dose [14]. We simulated T-DXd efficacy at a range of homogeneous expression levels from 10,000 receptors per cell (∼IHC1+ or low expression level seen in patients), to 40,000 receptors per cell using human PK parameters to determine the expression level at which we achieve tumor regression, i.e., tumor size below the starting volume at 21 days (**Fig 9**). As expected, at higher expression, we see a larger benefit from ADC dosing (smaller tumor volumes) because of the increase in number of ADC binding sites for direct payload delivery to cancer cells. We find that above 30,000 receptors per cell (approximately midway between ∼10,000 HER2/cell for IHC1+ and ∼100,000 HER2/cell for IHC2+), we start to see tumor volumes below the starting volume. (Note that IHC values are approximate, since levels vary across indications. CAPAN-1 may be considered high for pancreatic cancer cell lines [30] but low compared to breast cancer cell lines [20].) These results also align with clinical data that demonstrated that patients with higher IHC scores are more likely to benefit from treatment with T-DXd [31].

**Fig 9.**
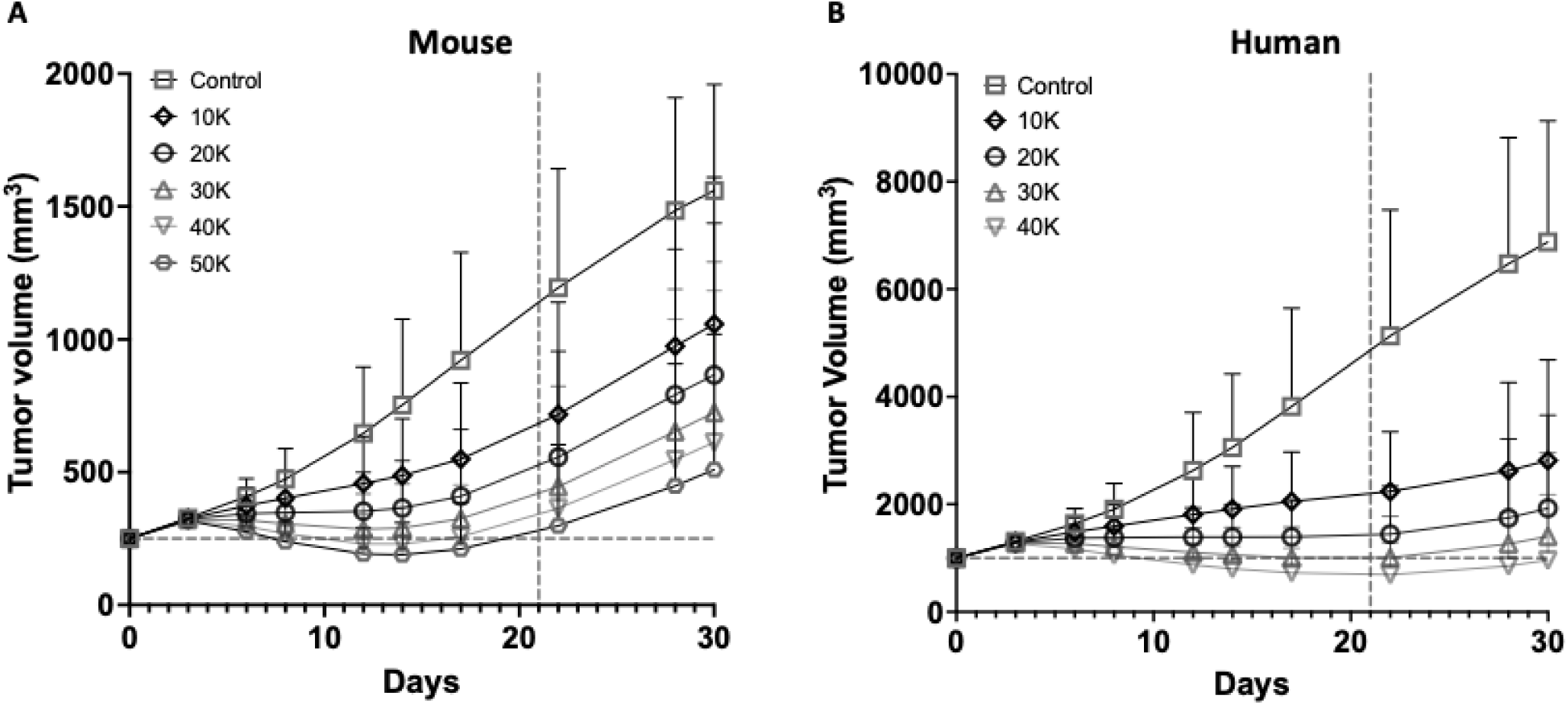
Minimum threshold of HER2 expression in mice and humans for efficacy of targeted delivery alone. The growth rates of tumors with varying low expression levels were simulated in *SimADC* following treatment with 5.4 mg/kg of T-DXd. Using tumor regression at 21 days as the threshold for efficacy, (A 50,000 HER2/cell were needed in mice while (B) 30,000 HER2/cell were sufficient in the human simulations. The difference occurs from scaling the pharmacokinetic parameters from mice to humans. Error bars represent standard deviation; each simulation point is the result of 300 simulation runs (100 simulations run in triplicate).

Further, because clinical data [32] have shown efficacy in patients with low or ultralow expression, we wanted to determine the mechanism(s) by which these patients may respond. With our calibrated and validated model, we next simulated T-DXd efficacy in low (10,000 HER2/cell) and negative (no HER2/cell) expression tumors with baseline and activated T cell killing, mac release, TME release, and systemic release (**Fig 2C**) to determine which mechanisms can contribute to tumor regression in patients.

### T cell activation is sufficient for strong responses in patients, with or without targeted delivery of T-DXd

Our simulations demonstrate that payload uptake from HER2 expression at or below the low (10,000 HER2/cell) level in patients is insufficient to drive tumor regression alone without additional mechanisms of action (**Fig 9**). We therefore next simulated tumor growth in patients with multiple mechanisms of action at play. We first tested the impact of T cells in a hypothetical low or negative expression tumor with clinically relevant PK parameters. After calibrating the immune effects of EphA2-PBD and EphA2-tubulysin, we used the model to examine the impact of targeted delivery versus baseline or activated T cell killing in HER2-low tumors (**Fig 2C**). A strength of the model is that we can calculate the individual contributions of DXd and T cell killing to efficacy. We simulated the efficacy of dosing the drug alone, with an additive relationship of payload and baseline T cell killing, and with a synergistic relationship incorporating the same level of T cell activation as seen in mice. We scaled the fraction of T cells in the tumor to 7.2% based on literature values reporting the number of CD8+ T cells/mm^2^ of tumor tissue for breast cancer patients to estimate the clinical scenario [33].

We find that tumor growth rates in HER2 low-expressing tumors with baseline T cell killing (only) are significantly reduced compared to T cell-negative tumors treated with T-DXd (**Fig 10A**). We see moderate efficacy with T-DXd drug alone, but this is not sufficient for tumor regression. However, the combination of baseline T cell killing and targeted delivery (additive) in low-expressing tumors is sufficient for tumor regression, generating good efficacy overall. With activated T cell killing and targeted delivery (synergistic), we see the best efficacy.

**Fig 10.**
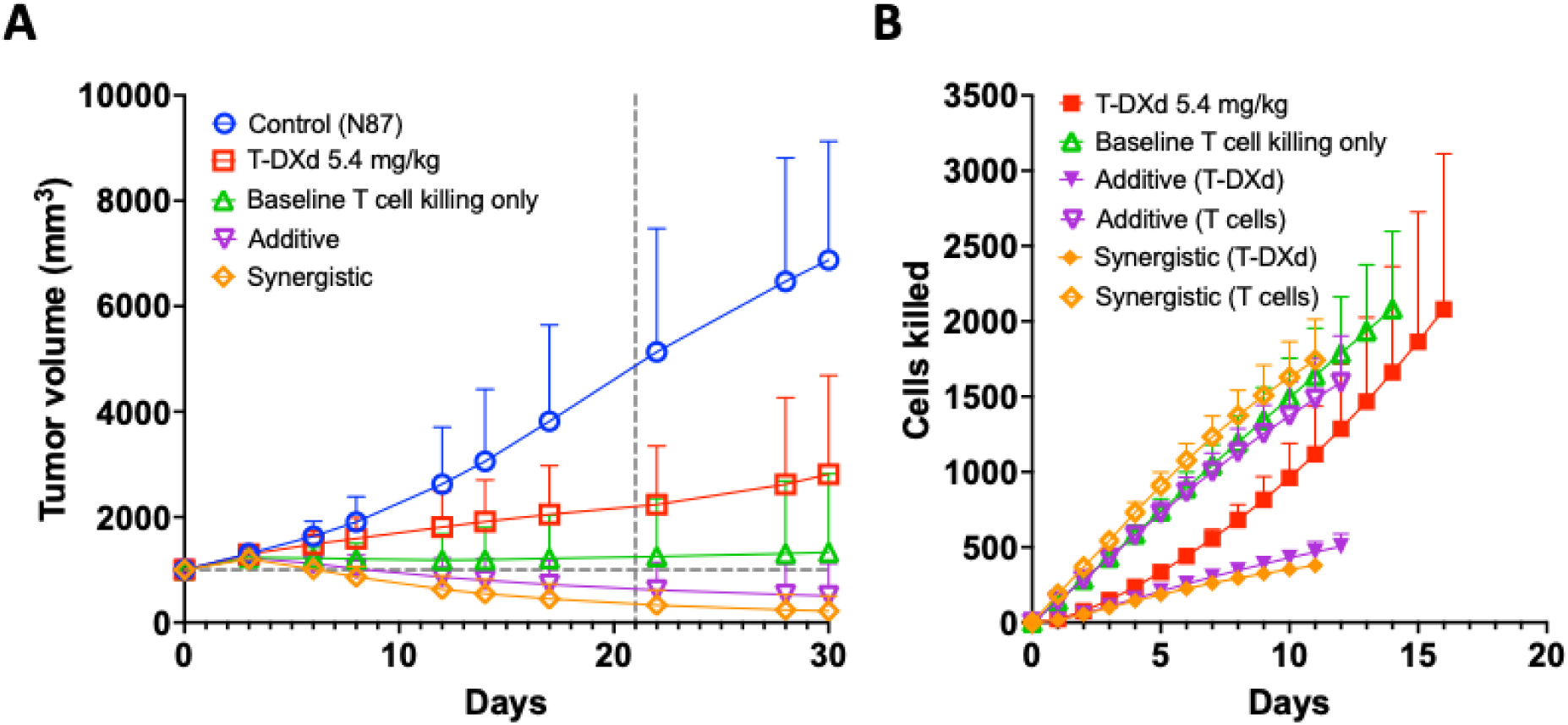
Predicted efficacy in HER2 low (10,000 HER2/cell) tumors in humans and cell killing mechanisms. Simulations tested control (no treatment), targeted delivery (5.4 mg/kg T-DXd only), baseline T cell killing only, and additive (baseline T cell killing plus targeted delivery) and synergistic (activated T cell killing plus targeted delivery) mechanisms. (A) With 10K HER2/cell, tumor growth is slowed but does not regress with targeted delivery of T-DXd alone. Infiltrating T cells without treatment also slow growth but don’t regress the tumor. However, a combination of 5.4 mg/kg of T-DXd with infiltrating T cells (baseline T cell killing) can regress the tumor, and activation of the T cells (synergy) can drive a strong response. (B) The first 2000 cells killed by each treatment regimen are plotted over time. Cells killed due to direct action of DXd (targeted delivery) vs T cell killing are plotted to separately to show their relative contributions. Compared to T-DXd treatment alone, the presence of T cells reduces the number of cancer cells killed by the drug with a further reduction following activation due to competing mechanisms. The T cell killing rate is not impacted as strongly by the T-DXd treatment, but it is substantially enhanced through activation. Error bars represent standard deviation; each simulation point is the result of 300 simulation runs (100 simulations run in triplicate).

To further investigate the interplay between ADC and T cell killing, we plotted the number of cancer cells killed per mechanism over time up to ∼2000 cancer cells killed (**Fig 10B**). For the additive relationship, we see that the number of cells killed by each mechanism decreases compared to their individual simulations, which occurs because the payload cannot kill a cell that has already been killed by a T cell and vice versa. However, we find that the first ∼2000 cancer cells are killed at an earlier timepoint with combined mechanisms than the individual mechanisms (12 days vs. 14 and 16 days). From these results, we conclude that for patients with more active intratumoral T cells, T-DXd can exert substantial efficacy even in low-expression tumors due to the synergistic relationship between T cells and ADCs. However, the presence or absence of these T cells is largely driven by the biology of the TME, and varying the intratumor T cell populations significantly impacts the overall efficacy (**Fig S6**). Therefore, the simulations primarily demonstrate the feasibility of the mechanism if a patient has significant tumor-infiltrating T cells.

### Mac release is sufficient for tumor regression

While our simulations show T cell killing with ADC treatment in low-expressing patients is sufficient for tumor regression, clinical results have so far failed to identify significant immune infiltration in responding patients [3], meaning other mechanisms may be at play in this population. We next predicted the impact of three target-independent mechanisms of payload release (mac release, TME release, and systemic release) on T-DXd efficacy in low and negative HER2-expressing tumors using human pharmacokinetic parameters and clinically relevant doses. We simulated each mechanism at the clinically approved dose of T-DXd (5.4 mg/kg) in tumors with NCI-N87 growth rates with negative (no receptors) and low (10,000 receptors per cell) expression using human PK parameters (**Fig 11**).

**Fig 11.**
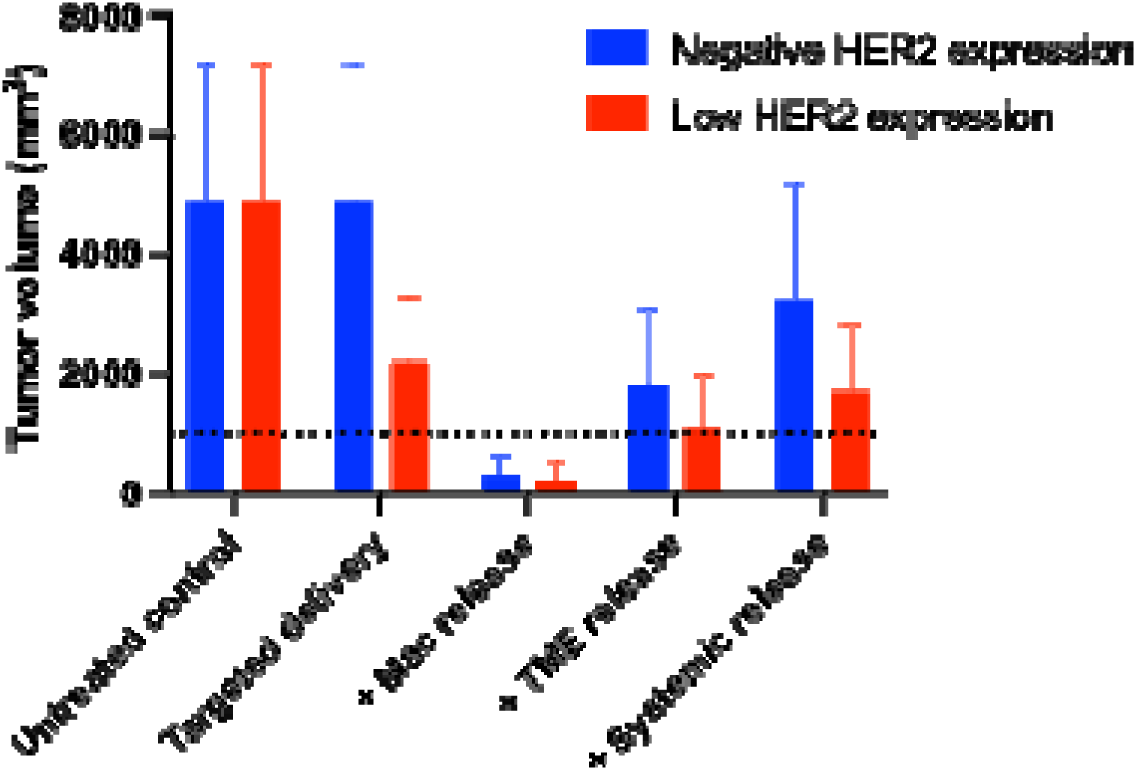
Target-independent mechanisms alongside targeted delivery show tumor growth control in humans. Simulated tumor volumes on day 21 (+/-standard deviation) were plotted for three target-independent mechanisms of action (mac release, TME release, and systemic release) acting in concert with targeted delivery of cancer cells. The dotted line shows initial tumor size. Lower HER2 targeting (10K HER2/cell) reduced tumor sizes to ∼50% with targeted delivery alone. Each of the three target-independent mechanisms added to this efficacy, with the greatest improvement from mac release, which was the only mechanism that resulted in tumor regression. Note that mac release and TME release are dependent on the number of macrophages and extracellular protease concentrations in the TME, respectively. Error bars represent standard deviation; each bar is the result of 300 simulation runs (100 simulations run in triplicate). The dotted line is the cutoff for 1000 mm^3^, in which tumor volumes below 1000 mm^3^ achieve regression.

As expected, we find that targeted delivery alone in negative expression tumors results in no efficacy. For low expression, there is low but significant efficacy relative to untreated tumors. With the efficient internalization of T-DXd, even 10,000 HER2/cell is sufficient to cut the growth rate in half (50% growth relative to an untreated control). However, this is not enough for tumor regression.

TME release showed moderate efficacy at 21 days and was able to result in stagnant growth when paired with low HER2 expression in the tumor. However, the greatest efficacy came from mac release with low or negative HER2 expression (**Figs 11 and S7**). We see significant tumor regression at 21 days, demonstrating the potential impact of large numbers of macrophages at clinically relevant doses, independent of direct HER2 targeting. Notably, the low expression tumors only have slightly greater efficacy than negatively expressing tumors because rapid macrophage internalization reduces the number of intact ADCs internalized by cancer cells compared to HER2 targeting alone.

To further investigate the mechanistic differences in response rates, we plotted the distribution of extracellular free payload and payload bound to intracellular targets, focusing on the TME release and mac release mechanisms. We examined the distribution at 8 hours, 1 day, and 5 days. We found that at early times, TME release allows for rapid extracellular payload release and tumor uptake, but the payload starts to clear and is almost completely washed out of the tumor by day 5 (**Fig S8**). For mac release, we see some extracellular free payload but little uptake in the first 8 hours (primarily due to slower macrophage payload release). However, high concentrations of payload are released by day 1, followed by high cellular uptake and payload binding. At day 5, high concentrations of extracellular free payload and high tumor uptake are maintained, indicating that the macrophages can provide sustained release in the tumor, leading to greater efficacy.

### Systemic release is not sufficient for tumor regression

When the contribution of systemic release and tumor uptake is included, tumor growth is reduced 35% from an untreated HER2 negative tumor. (**Fig 11**) However, the growth reduction is lower than direct ADC efficacy in low HER2 expression tumors or other target-independent mechanisms of payload release. While the payload concentration in the plasma is above the *in vitro* IC_50_ value, blood flow limitations and tissue partitioning result in lower free drug concentrations at the site of action in the tumor, limiting efficacy. This is also consistent with low clinical response rates with continuous intravenous infusions of exatecan, as demonstrated with simulations of slow infusion (**Fig S9**).

To further investigate the impact of systemic release on ADC efficacy, we used the model to simulate high-expressing tumors in mice and humans dosed with T-DXd and T-MMAE over time. Previous literature raised the point that this low persistent exposure to free payload from ADC dosing could drive efficacy [13]. Since *SimADC* can now simulate efficacy from systemic release, we matched the pharmacokinetics for free payload in the blood after dosing T-DXd and T-MMAE in both mouse and human models and simulated the equivalent dose to determine the contribution of systemic release on efficacy. Systemic release after dosing ADC results in higher and more sustained concentrations in humans because of the constant dissociation from the ADC and slower payload clearance. For T-DXd free payload, we saw negligible impact in mice as expected due to rapid payload clearance. The impact was larger in humans but not enough for tumor regression, especially when compared to targeted delivery (**Fig S10**). For T-MMAE, we saw little contribution from systemic release when compared to the dose used in the associated pharmacokinetic curve (10 mg/kg) (**Fig S11**).

### Targeted delivery is the dominant driver of ADC efficacy with varying contributions from target-independent effects

To understand how the three target-independent mechanisms tested may work together, we ran a series of simulations to determine if combinations of mechanisms can achieve tumor regression or if there is an antagonistic effect from multiple mechanisms working at the same time (**Figs 12 and S12)**. The addition of systemic release to TME release is capable of achieving tumor regression, which was not previously achieved by TME release alone, and the addition of systemic release to mac release further drives tumor regression. Additionally, we did not see antagonistic effects for any combination mechanism. Overall, however, targeted delivery with moderate and high HER2-expressing cells yields the strongest response.

**Fig 12.**
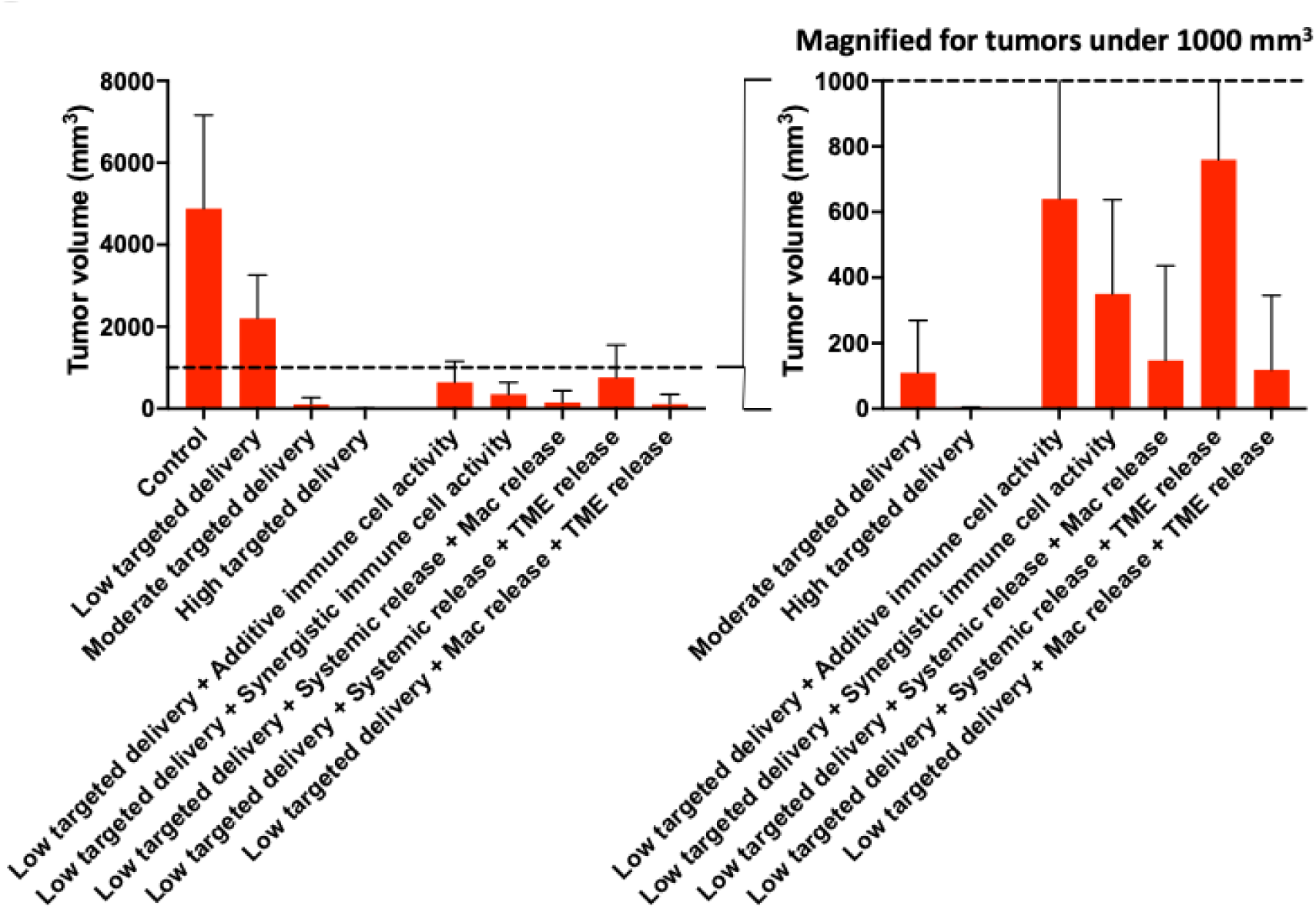
Efficacy of combined mechanisms alongside targeted delivery to moderate and high HER2-expressing tumors. The addition of systemic release to TME release was enough to regress tumors. The addition of either systemic release or TME release to mac release was enough to drive further tumor regression. For comparison with the combined target-independent mechanisms, tumors with moderate HER2 expression (100,000 HER2/cell) and high HER2 expression (1,000,000 HER2/cell) underwent simulated treatment, and both resulted in very strong efficacy, consistent with the high efficiency of targeted payload delivery. Error bars represent standard deviation; each bar is the result of 300 simulation runs (100 simulations run in triplicate). The dotted line is the cutoff for 1000 mm^3^, in which tumor volumes below 1000 mm^3^ achieve regression.

To summarize all the individual mechanisms alone and in combination, **Table 1** highlights which results show high efficacy (tumor regression), moderate efficacy (∼50% reduction in tumor growth) or low efficacy (< 40% reduction in tumor growth). Overall, high or moderate HER2 expression, immune cell activation, or mac release were the strongest drivers of tumor regression.

**Table 1.** Summary of efficacy across varying HER2 expression and target-independent mechanisms.

|  | Targeted delivery<br>(Expression level*) | T cell activation | Mac release | TME release | Systemic release | Efficacy |
| --- | --- | --- | --- | --- | --- | --- |
| Targeted<br>delivery only | +++ |  |  |  |  | High |
|  | ++ |  |  |  |  | High |
|  | + |  |  |  |  | Moderate |
|  | - |  |  |  |  | Low |
| Non-target<br>mechanisms | + / - | ✓ |  |  |  | High |
|  | + / - |  | ✓ |  |  | High |
|  | + / - |  |  | ✓ |  | Moderate |
|  | + / - |  |  |  | ✓ | Moderate/Low |
| Combined<br>mechanisms | + / - |  | ✓ | ✓ |  | High |
|  | + / - |  | ✓ |  | ✓ | High |
|  | + / - |  |  | ✓ | ✓ | Moderate |

\* +++ is ~1,000,000 HER2/cell (IHC3+), ++ is ~100,000 (IHC2+), + is ~10,000 (IHC1+), and negative
✓ indicates active mechanism
+ / - represents simulations that were run for both low (+) and negative (-) HER2 expression

## Discussion

ADCs are complex therapeutics with multiple molecular interactions, both within the tumor microenvironment and systemically (**Fig. 1**, [34]). These interactions can be studied in mice, but the magnitude of the effects can differ when scaling to human clinical doses and pharmacokinetics. Further, many clinical payloads have low efficacy in the mouse cancer cells used for immunocompetent models, resulting in stronger apparent immune effects versus payload effects in these models. Conversely, human cancer cells used in immunodeficient xenograft models have high payload effects but minimal immune effects. Humanized mouse models include human immune cells and cancer cells, but many of these show reduced myeloid cell infiltration, which can play a significant role in payload release [35]. These factors limit the use of preclinical models to compare multiple mechanisms of ADC action. Clinical data provide the most direct insight into mechanisms, but clinical data are limited. Here, we use a computational approach with our validated computational model, *SimADC*, to integrate data from multiple preclinical and clinical studies and provide a framework for analyzing and understanding clinical data (**Fig. 2**). We compared the impact of T cell activation, free payload in the plasma, macrophage uptake, and extracellular cleavage on T-DXd efficacy in low and negative HER2 expression systems, to quantitatively compare the mechanisms driving the success of T-DXd at these expression levels.

### Immune cell interactions via ADC activation can enhance tumor regression

Clinical data demonstrate that immune cells like tumor-infiltrating lymphocytes (TILs), particularly CD8+ T cells, are associated with better ADC treatment responses for HER2-positive cancer [36]. Additionally, ADCs are being paired with checkpoint inhibitors in the clinic, including the FDA-approval of enfortumab vedotin in combination with pembrolizumab and positive clinical results with sacituzumab govitecan and pembrolizumab. Our simulations demonstrate how activated T cells from ADC treatment can contribute to further tumor regression compared to the baseline T cell killing. Importantly, the model was only able to capture the in vivo effects when the efficiency of T cell killing was increased, confirming this is driven by T cell activation and not just independent additivity of ADC efficacy with baseline T cell activity (**Figs 3 and 4**). This activation could be driven by damage-associated molecular patterns (DAMPS) from immunogenic cell death [16] or direct effects of the payload on immune cells that can drive stimulation [9,16].

These results are consistent with some clinical observations [37], but in the case of HER2 low/negative tumors, many patients do not show clear immune responses. In our simulations, the fraction of T cells in the tumor are based on clinical data showing an average of 360 CD8+ T cells/mm^2^, but many patients have fewer T cells [33]. The percentage of T cells in the tumor and activation status can greatly influence the simulated response to ADC treatment, as demonstrated in **Fig S6**. The significant impact of activated T cells is why many immune-based therapies focus on driving strong immune responses with CD8+ T cells (i.e. T cell engagers, CAR T cell therapy) [38]. However, there are patients who don’t respond to immune-stimulating agents or have fewer infiltrating T cells that don’t benefit from such treatments but still show efficacy from ADC therapy [3]. Hence, we investigated mechanisms based on just the free payload of the ADC to determine why patients may still respond despite low or no HER2 expression.

### Target-independent mechanisms of ADC payload release

We calibrated *SimADC* to quantify the efficacy from three different mechanisms of target-independent free payload release and determine if they can drive tumor regression when scaled to clinical doses and pharmacokinetics. These mechanisms include mac release, TME release, and systemic release (**Fig 2C**).

#### Macrophage payload release can drive tumor efficacy

Intratumoral macrophages encompass a range of cellular phenotypes that can have beneficial effects, such as antibody-dependent cellular cytotoxicity (ADCC) and antibody-dependent cellular phagocytosis (ADCP), or immunosuppressive effects, embodied in the broad category of myeloid derived suppressor cells (MDSCs). In addition to immune effects, intratumoral macrophages can also release ADC payloads in the tumor. These processes are often activated through Fc binding, in which the Fc region of the antibody binds to Fc receptors on macrophages. Li et al. [10] demonstrated how the presence of macrophages drives ADC efficacy in some models, where even the mac release from isotype control was sufficient for efficacy. In our simulations with T-DXd, we see strong efficacy from this mechanism (**Fig 6**), indicating it has the potential to drive T-DXd efficacy at low and negative expression levels (**Fig 11**). Importantly, the number of myeloid cells and Fc receptor expression both impact the magnitude of this mechanism (**Fig S5**), so additional work on quantifying the range in expression across different patient populations would improve the predictions. Overall, this mechanism showed the highest responses for non-targeted delivery, capable of exceeding the response from low target expression and approaching that of moderate expression (although less effective than high HER2 expression).

#### Sustaining free payload in the tumor can impact efficacy

In our simulations, TME release resulted in moderate efficacy compared to mac release over 21 days (**Fig 11**). Like mac release, this mechanism is also sensitive to the specific rate within the tumor. The rate of cleavage was calibrated to experimental data in a mouse model (**Fig 7**), though this value could vary based on the number of cleavage proteases released by other cells in the tumor, including cancer cells or macrophages. The cleavage rate depends on the specific linker and environmental factors (e.g. number of proteases, pH, or signaling pathways). The sensitivity can also be seen across mouse models. For example, the same tumor xenograft is reported to show significant extracellular cleavage and efficacy when grown in SCID mice, while it shows negligible efficacy when grown in nude mice [11,20]. Clinically, T-DXd shows greater efficacy in HER2 low or negative breast cancer tumors than other cancer indications [37], which could be related to differences in the TME and non-target mediated mechanisms. More work on understanding the clinical role of extracellular proteases and macrophage/myeloid cell uptake and release is needed.

Comparing mac release with TME release, we observed differences in the timescales at which payload is taken up into cancer cells (**Fig S8**). For TME release, the rapid extracellular cleavage resulted in a large amount of free payload in the extracellular space after 8 hours, followed by clearance of the free payload through day 5. Without any binding of the ADC to tumor cells or macrophages, the extracellular concentrations are largely driven by enhanced permeability and retention (EPR) effects [39], lowering the concentration of ADC susceptible to payload cleavage. Methods of anchoring the drug-conjugate in the TME, such as binding slow or non-internalizing targets, could improve this mechanism [12]. In contrast, mac release has an initial delay while the Fc-internalized ADC released the payload that then must diffuse out of the macrophage. However, the intracellularly-released free payload diffuses into the extracellular space at a slower but more sustained rate. By day 5, we see higher payload bound in the tumor compared to extracellular payload, indicating that macrophages can prolong the presence of free payload in the tumor through a slower release.

TME release and mac release are not mutually exclusive, and we see slightly more tumor regression when both mechanisms are active. Additionally, extracellular proteases are released by a variety of cell types within a tumor, including macrophages, such that the contribution of macrophages could include both intracellular and extracellular cleavage. However, the ADC structure can impact these mechanisms independently. Fc-engineering (e.g. Fc-silent ADCs) could reduce mac release while more stable linker chemistry could reduce TME release. Likewise, the choice of DAR would have differential impacts on these mechanisms, with a saturating dose of a low DAR ADC having less mac release than a saturating dose of high DAR ADC, while TME release would be more dependent on the total payload dose independent of DAR. Therefore, both mechanisms have important implications in ADC design.

#### Systemic release

It has been observed that the concentration of systemic release is above the *in vitro* IC_50_, indicating it could potentially contribute to ADC efficacy [13]. However, even small molecules have delivery limitations in tumors, where the concentration at the site of action may not equilibrate with the plasma due to reduced perfusion from collapsed blood vessels and poor flow from elevated interstitial pressure, among other factors. To quantify the clinical impact of free payload from T-DXd, we calibrated *SimADC* to simulate efficacy from clinical concentrations of free payload in the blood (**Figs 8, S10, and S11**). Systemic release after dosing an ADC results in higher and more sustained concentrations in humans because of the constant dissociation from the ADC and slower payload clearance, and the simulated efficacy of free payload is larger in humans than in mice. However, the simulations indicate it is still smaller than other target-independent mechanisms of action (**Fig 11**).

To further test this result, we simulated and compared efficacy for CIVI dosing of free exatecan over 21 days (**Fig S9**). Unlike the high bolus dosing of free drug (‘payload’) from classic chemotherapy treatment, the sustained low concentration of drug did not result in sufficient uptake for tumor regression. These simulation results are consistent with clinical studies of continuous infusions or repeat dosing aimed at achieving sustained levels of exatecan [22,40,41]. Likewise, approved ADCs utilizing MMAE as a payload have similar C_max_ values and actually lower AUC values of free MMAE exposure than discontinued ADCs, implying the free payload exposure isn’t the main contributor to efficacy (**Fig S13**) [42]. This has important implications for the design of new agents, where reducing systemic levels of free payload could reduce toxicity without impacting efficacy. This generally can’t be achieved solely through improved linker stability, however, since intact ADC uptake, intracellular release in healthy tissue, and diffusion back into the blood also contributes to toxicity. Overall, these results support the importance of the role of antibodies in delivering payload into the tumors rather than systemic payload release.

### Implications for T-DXd efficacy in HER2 low and negative tumors

Our *SimADC* framework can be applied to any ADC, and the initial training utilized data from multiple ADCs and small molecules. However, the focus of the simulations with multiple mechanisms in this work was on treatment with T-DXd, and the results are largely consistent with clinical data on T-DXd. First, Mosele et al. [3] showed that T-DXd responses correlate with HER2 expression with the highest efficacy for HER2+, then HER2-low, and finally HER2-null patients. These results align with the importance of targeted payload delivery into cancer cells in *SimADC* (**Fig 12)**. The non-target mediated mechanisms are most important at low expression ranges, where insufficient HER2-targeted delivery exists to drive responses. For the non-target mediated mechanisms, the impact of immune cell killing and activation by ADCs in the simulations is sufficient to drive strong responses when the preclinical response rates are scaled to the clinic. Kapil et al. [43] showed stronger progression free survival in HER2+ patients with higher levels of TILs [37]. This implicates immune effects in the outcome of these breast cancer patients. However, TILs did not stratify patients with low or negative HER2 expression, and similarly, Mosele et al. [3] did not detect significant changes in PD-L1 expression or immune cells while on treatment for patients with HER2-low or non-expressing tumors. Therefore, while T cell immune effects can play a role when TILs are present, additional target-independent mechanisms may play a significant role in other contexts.

The efficient internalization of HER2 means that even very low levels of HER2 expression (∼10,000 HER2/cell) can significantly contribute to efficacy. These levels are similar to the HER2 expression reported in healthy breast tissue [44,45]. With typical human cancer cell sensitivity to DXd, *SimADC* did not show tumor regression with 10,000 HER2/cell, but levels as low as 30,000 HER2/cell were sufficient to drive tumor regression, dramatically below the 1,000,000+ HER2/cell that can be seen in overexpressed tumors. Still, there are many examples of responses in patients with undetectable HER2 expression, indicating target-independent mechanisms may play a role. Interestingly, Eisses et al. [46] shows significant trastuzumab uptake in IHC 0, 1+, and 2+ patients, implying efficient delivery and concentration within the tumor microenvironment across a range of expression levels. This could be driven by non-specific vascular effects (EPR effect) and/or binding to Fc receptors within the tissue. This tumor uptake would be consistent with TME or mac payload release. These results call for a better understanding of the TME and stromal cells in clinical cancers.

Despite the incorporation of multiple mechanisms of action in a single uniform framework, *SimADC* has limitations. First, the complex biology controlling the number and activation of immune cells within the TME is not captured in a predictive fashion. Rather, the simulations here are trained on syngeneic models to capture the magnitude of these effects, so the T cell killing is imposed rather than an emergent effect. Additional features, such as macrophage and T cell crosstalk, T cell recruitment, checkpoint suppression, and T cell exhaustion, can be added to the model in future work as more quantitative biological information becomes available [47]. Likewise, the impact of other immune cell types like ADCC from NK cells and ADCP from macrophages are not included in the current model. This choice was based on the dominance of T cell killing in the syngeneic models used to train *SimADC*, but these could be added later. These mechanisms may play a more substantial role in disseminated metastatic disease than large established tumors. Similarly, the effects of target-independent payload release were trained on preclinical models; tumors with few Fc-receptor expressing immune cells or extracellular proteases would have lower responses from these mechanisms. The systemic metabolism of ADCs from non-specific uptake or targeted internalization in healthy tissue and systemic deconjugation of payload can significantly alter the systemic free payload. For example, the plasma concentrations of sacituzumab govitecan are much higher and more transient than T-DXd [48]. Another important consideration is HER2 expression on cancer cells or stem-like cells [49,50]. Killing of HER2-positive stem-like cancer cells falls under the mechanism of direct targeting for ADC cell death, but the magnitude of the impact would be greater than the number of HER2-positive cells given the larger potential role in tumor growth. Despite these limitations, SimADC provides a useful tool to generate hypotheses, interpret preclinical and clinical results, and design next-generation targeted agents to maximize the efficacy of ADCs by leveraging data from multiple sources (*in vitro*, *in vivo* models (mouse, rat, NHP), and clinical data).

In conclusion, the incorporation of immune cells and target-independent mechanisms of payload release into *SimADC* enables efficacy predictions of ADCs in the clinic by scaling multiple mechanisms of action from in vitro, in vivo, and clinical data. It is possible to input patient-specific parameters such as payload sensitivity, immune cell counts, and immune cell activation and predict the efficacy of ADCs with different payload, linker, and antibody designs with multiple dosing strategies. Tools like these can also reduce the number of animal experiments and aid in clinical trials, potentially preventing toxic outcomes for cancer patients. We can also use such simulation tools to determine optimal PK parameters for novel ADCs and test the impact and sensitivity of parameters like binding affinity, internalization rates, and drug diffusion on different types of cancer. By incorporating immune cells, we can predict the effects of ADCs on activating a patient’s immune system to kill cancer cells, thereby leveraging multiple mechanisms of action. This could be applied to other payload classes, such as ISACs (immune stimulatory antibody conjugates) and offset the potential toxicity to patients during development. Therefore, computational tools like *SimADC* can aid in experimental design, data interpretation of preclinical and clinical results, and novel therapeutic design to improve ADC efficacy.

## Methods

### SimADC model design

*SimADC* is a hybrid agent-based model built to study ADCs at multiple spatiotemporal scales. As originally described in Menezes et al. [51,52], the model simulates a 2D slice from a 3D tumor and assumes the ADC concentration along the blood vessel to be approximately constant, since ADC diffusion is limited by its extravasation through the vessel wall. We also assume that the drug dynamics on the grid are representative of the entire tumor, and that the tumor volume is proportional to the number of cells in the simulation. When ADC is administered, the drug diffuses onto the grid through the blood vessels, calculated by PDEs, and the individual cell concentrations are calculated using ODEs for binding, internalization, degradation, and payload release. When a cell dies, the cell remains on the grid for 2.5 days (calibrated from experimental mouse data) before it is removed, to represent the time it takes for a cell to be removed from the tumor. When dead cells are removed or a cell divides, the agents shuffle based on an algorithm implemented to maintain a circular tumor shape.

The model is stochastic, using probabilities to reflect biological heterogeneity. The tumor growth rate, which is represented by a cancer cell doubling time, and the fraction of active blood vessels are chosen from a pseudo-normal distribution using Latin Hypercube Sampling (LHS) [53]. Blood vessels are also placed randomly on the grid to capture the heterogeneity in vessel distribution. Cancer cells die based on the concentration of payload bound to its target using Michaelis-Menten kinetics, in which the concentration determines the probability of cell death for individual cells. A random number is generated at every time step and compared to this probability to determine whether or not a cell will die. All model variables, equations and baseline parameters are listed in **Equations S1** and **Table S1**.

### Payload dosing in nude and syngeneic mice

The tumor microenvironment (TME) is a complex network of malignant cells, immune cells, tumor vasculature, and other stromal cells. The immune system plays a major role in the TME, contributing to both tumor growth and tumor cell death. CD8+ T cells are the main effectors of cell-mediated adaptive immune responses, as they can directly kill cancer cells and infected cells. To capture the functionality of CD8+ T cells, we established the following rules:

1. T cell steady state: T cells appear on the whole grid at initialization and remain for the entire simulation. As the mass of cancer cells expands during tumor growth, the boundary of the tumor will encompass more and more T cells, representing recruitment (and vice versa, as the tumor shrinks), keeping a constant density of ∼10%, based on literature values [8,54]
2. T cell randomization: T cells appear randomly on the grid since we assume sufficient vascularization throughout the tumor.
3. T cell killing: T cells kill neighboring cancer cells one at a time with a specified probability.

The parameter involved in T cell killing is pKill, the probability that a T cell will kill one of its cancer cell neighbors. T cells check their surrounding grid positions for cancer cell neighbors and select one at random. Then a random number between 0 and 1 is generated and the cancer cell dies if the random number generated is below pKill. Similar to how cancer cells die via ADC, the cell stays on the grid for 2.5 days before it is removed from the grid to represent removal of the dead cell/debris.

T cell agents are split into two populations, active and inactive, each with their own probability of killing neighboring cancer cells (the inactive population having a much smaller, intrinsic probability of cell killing). These killing parameters, along with our model PD parameters, were calibrated to literature data from Rios-Doria et al. [8], who reported efficacy curves for both immunodeficient and immunocompetent CT26 mouse models dosed with multiple doses of EphA2-Tubulysin and EphA2-PBD. The percentage of active T cells for the control, 5 mg/kg of EphA2-Tubulysin, 0.3 mg/kg of EphA2-PBD was reported and used for model calibration.

The tumor growth rate for the CT26 cell line was calibrated in the immunodeficient model to isolate payload killing from immune cell killing and study each mechanism both separately and additively. CT26 is commonly used mouse cell line for immunogenic tumors [55]. Therefore, this captures the upper end of T-cell efficacy for comparison with no T-cell simulations. Next, we fit our model pharmacodynamic parameters to the immunodeficient mouse model efficacy studies for EphA2-Tubulysin and EphA2-PBD. These pharmacodynamic parameters are the Michaelis-Menten rate constant, K_m_, and the probability of cell killing, P_max_. We used antibody pharmacokinetic parameters from internal data [56] for the expression, binding and internalization rates, and clearance for the EphA2 antibody, and we used payload pharmacokinetic parameters from previous work for PBD and MMAE (used for Tubulysin).

Next, the baseline immune effects of T cells had to be captured in the absence of ADC treatment. The P_kill_ parameter established the baseline T cell killing probability for a T cell to kill and adjacent cancer cell. Once the tumor T cell density was set equal to the experimentally measured value, P_kill_ was calibrated to tumor growth studies to account for the difference in growth rates for the same cell line grown in an immunodeficient versus an immunocompetent mouse strain from Rios-Doria et al. [8] experiments. A baseline T cell killing probability was able to capture the slower growth rate of tumors with T cells in the syngeneic model versus the growth rate of these cells in an immunodeficient or a CD8 T cell depleted model (**Fig 3**).

### Macrophages for payload release

Macrophages in *SimADC* are placed randomly on the simulation grid, similar to T cells and cancer cells. To study the impact of Fc binding on ADC efficacy, the macrophage agents utilize the same equations used for the cancer cells that represent ADC and antibody binding and internalization. This binding to the macrophages represents Fc receptor binding to the Fc region of the antibody. The pharmacokinetic parameter values (binding, internalization, and clearance kinetics) for macrophages and cancer cells were estimated from literature values and/or internal lab data [19,57].The percentage of macrophages in the tumor was estimated from IHC analysis in L-428 tumors showing high macrophage infiltration [10]. We assume that the number of macrophages being recruited to the tumor and dying within the tumor are equal so that the percentage of macrophages on the grid stays constant. Because of increased or sustained levels of macrophages seen in literature after ADC dosing [3,10], we also assume that the macrophages are not affected by the drug (i.e. killed) apart from binding, internalization, and payload release.

We first fit the tumor growth rates to the control for L428 tumors from the *in vivo* efficacy data [10]. We then used pharmacokinetic parameters for IgG from literature [17,18] for cancer cell uptake and fit our MMAE pharmacodynamic parameters to the *in vivo* efficacy curves for each ADC. For our macrophage parameters, we used internal data measuring the rates of Fc binding, internalization, and receptor expression (387,500), and used tumor staining images from Li et al. [10] to determine the macrophage density on the simulation grid (50%). To reduce computational run times, we halved the density of macrophages on the grid and doubled the receptor expression (25% macrophages with 775,000 receptors per cell) to emulate the same amount of ADC uptake. For both simulations, to calculate intratumoral payload, we summed up the average intracellular, extracellular, and target-bound payload over time for comparison, along with our simulated tumor volumes (**Fig 6**).

### Extracellular cleavage for TME release

Our cleavage parameter k_Cleave_ was fit to *in vivo* animal data from Tsao et al. [11] that demonstrated efficacy with T-DXd in a negative expression cell line, MDA-MB-468. We first fit the tumor growth rates to the untreated curve. We then set the model target expression to zero and simulated our previously-used T-DXd PKPD parameter set at the specified dose (10 mg/kg, weekly). We tuned k_Cleave_ to fit two sets of efficacy data in the same cell line, and used this parameter to simulate T-DXd in our simulated NCI-N87 low and negative expression cell lines.

### Fitting free payload pharmacokinetics

We fit the model clearance parameters to multiple pharmacokinetic curves to model dosing free payload to the tumor. We used literature data from sources [13,20,26,29,58] to fit the pharmacokinetics of free payload after dosing free payload alone and after ADC administration, for multiple types of payloads, and from both mouse models and clinical patient data. To fit each clearance curve, we used a nonlinear regression two phase decay model in GraphPad Prism to fit the experimental data and obtain the constants for the alpha and beta clearance phases. We then calculated the clearance rates for the main and peripheral model compartments to use in simulation (**Table 2**).

**Table 2.** Pharmacokinetic parameters for plasma concentration of free payload in mice and humans.

| Symbol | Parameter | Mouse DXd [20] | Human DXd [13] | Mouse MMAE [29] | Human MMAE [13] | Unit |
| --- | --- | --- | --- | --- | --- | --- |
| CL<sub>10</sub> | Total antibody clearance | $2.21 \times 10^{-6}$ | $1.86 \times 10^{-6}$ | $2.00 \times 10^{-5}$ | $2.03 \times 10^{-6}$ | s<sup>-1</sup> |
| CL<sub>12</sub> | Antibody clearance for compartment 1 to 2 | - | $4.98 \times 10^{-7}$ | $8.06 \times 10^{-6}$ | - | s<sup>-1</sup> |
| CL<sub>21</sub> | Antibody clearance for compartment 2 to 1 | - | $5.11 \times 10^{-7}$ | $8.00 \times 10^{-6}$ | $6.77 \times 10^{-6}$ | s<sup>-1</sup> |

For the experimental data, the concentration at day 0 is typically measured immediately after dosing, as free payload undergoes rapid initial distribution and clearance. In *SimADC*, one timestep is equivalent to 10 minutes, such that a lot of small molecule payload has redistributed and may already be cleared. To account for this change, we used an equivalent dose to match the initial concentration of drug in the blood for each clearance curve. The pharmacokinetic parameters for free DXd and MMAE were obtained from Khera et al. [6]

For the CIVI simulations, we validated our free payload PK parameter set to estimate the plasma concentration curve based on the dose reported in Garrison et al. [22] Similar to the previous PK curve fittings, we found a dose that more closely matched the magnitude of plasma concentration from the experimental data to use as our equivalent model dose. We then doubled the maximum probability of cell killing to match the difference in potency between exatecan and DXd and simulated this dose in dosing regimens of 10 and 21 continuous days of dosing as outlined in Garrison et al.

### Scale up from mouse to human parameters

The pharmacodynamic parameters for T-DXd were fit to internal efficacy mouse data using NCI-N87 tumors, as well as *in vitro* data reported in Ogitani et al.[21]. After calibrating each payload release mechanism to mouse data, we scaled up our model to human parameters by changing the host weight, plasma volume, clearance rates, and starting tumor volume in the model (**Table 3**). The tumor growth rates and pharmacokinetic and pharmacodynamic parameters used for fitting mice data were kept the same.

**Table 3.** Pharmacokinetic and simulation parameters of ADCs in mice and humans.

| Symbol | Parameter | Mouse | Human | Unit |
| --- | --- | --- | --- | --- |
| BW | Host weight | 0.025 | 70 | kg |
| $V_1$ | Central compartment volume | 0.0012 | 3 | L |
| $CL_{10}$ | Total antibody clearance | $4.55 \times 10^{-6}$ | $1.858 \times 10^{-6}$ | $s^{-1}$ |
| $CL_{12}$ | Antibody clearance for compartment 1 to 2 | $8.06 \times 10^{-6}$ | $4.982 \times 10^{-7}$ | $s^{-1}$ |
| $CL_{21}$ | Antibody clearance for compartment 2 to 1 | $1.33 \times 10^{-5}$ | $5.109 \times 10^{-7}$ | $s^{-1}$ |
| $V_0$ | Starting tumor volume | 250 | 1000 | $mm^3$ |

