## Supplemental data for "Agent-based modeling demonstrates how target-independent processes supplement killing by antibody-drug conjugates in cancer therapy"

Supplemental Material

Melissa C. Calopiz^1^, Jennifer J. Linderman^1,2^, Greg M. Thurber^1,2,3*^

^1^Department of Chemical Engineering, University of Michigan, Ann Arbor, Michigan, United States of America

^2^Department of Biomedical Engineering, University of Michigan, Ann Arbor, Michigan, United States of America

^3^Rogel Cancer Center, University of Michigan, Ann Arbor, Michigan, United States of America

* Corresponding author

 (GMT)

**Equations S1. *SimADC* model variables, equations, and parameters**

From Menezes et al. 2020 [1] and Menezes et al. 2022 [2]

**Drug Dynamics Variables**

| **Variable** | **Unit** | **Description** |
| --- | --- | --- |
| ADC_c_ | nM | ADC in the central compartment |
| ADC_p_ | nM | ADC in the peripheral compartment |
| Ab_c_ | nM | Unconjugated antibody in the central compartment |
| Ab_p_ | nM | Unconjugated antibody in the peripheral compartment |
| ADC | nM | Free ADC in the tumor |
| Ab_free_ | nM | Free unconjugated antibody |
| T_free_ | nM | Free target |
| B_ADC_ | nM | ADC bound to target |
| B_ADC,lys_ | nM | ADC bound in the lysosome |
| P_lys_ | nM | Payload in the lysosome |
| P_int_ | nM | Free intracellular payload |
| P_b_ | nM | Bound payload to intracellular target |
| P_ext_ | nM | Free extracellular payload |
| B_Ab_ | nM | Bound unconjugated antibody |

**ADC Plasma Dynamics**

$$\left( 1 \right) \frac{d\left[ {ADC}_{c} \right]}{dt}=-k_{10}\left[ ADC \right]-k_{12}\left[ {ADC}_{c} \right]+k_{21}[{ADC}_{p}]$$

$$\left( 2 \right) \frac{d\left[ ADC_{p} \right]}{dt}=-k_{21}\left[ {ADC}_{p} \right]+k_{12}\left[ ADC_{c} \right]$$

$$\left( 3 \right) \frac{d\left[ {Ab}_{c} \right]}{dt}=-k_{10}\left[ {Ab}_{free} \right]-k_{12}\left[ {Ab}_{c} \right]+k_{21}[{Ab}_{p}]$$

$$\left( 4 \right) \frac{d\left[ {Ab}_{p} \right]}{dt}=-k_{21}\left[ {Ab}_{p} \right]+k_{12}\left[ {Ab}_{c} \right]$$

**Trastuzumab and ADC Extravasation and Diffusion**

$$\left( 5 \right) -D_{eff}\frac{d\left[ ADC \right]}{dx}=P(\left[ ADC \right]_{plasma}-\frac{\left[ ADC \right]}{\varepsilon})$$

$$\left( 6 \right) -D_{eff}\frac{d\left[ {Ab}_{free} \right]}{dx}=P(\left[ {Ab}_{free} \right]_{plasma}-\frac{\left[ {Ab}_{free} \right]}{\varepsilon})$$

$$\left( 7 \right) -D_{eff}\frac{d\left[ P_{ext} \right]}{dx}=P(\left[ P_{ext} \right]_{plasma}-\frac{\left[ P_{ext} \right]}{\varepsilon_{p}})$$

$$\left( 8 \right) \frac{\partial C}{\partial t}=D(\frac{\partial^{2}C}{\partial x^{2}}+\frac{\partial^{2}C}{\partial y^{2}})$$

Diffusible species (e.g. [ADC], [Ab_free_], [P_ext_]) incorporate a separate diffusion step, equation (8).

**Trastuzumab and ADC Drug Dynamics**

$$\left( 9 \right) \frac{d\left[ ADC \right]}{dt}=-k_{on}\frac{\left[ ADC \right]}{\varepsilon}\left[ T_{free} \right]+k_{off}\left[ B_{ADC} \right]$$

$$\left( 10 \right) \frac{d\left[ T_{free} \right]}{dt}=R_{s}-k_{on}\frac{\left[ ADC \right]}{\varepsilon}\left[ T_{free} \right]+k_{off}\left[ B_{ADC} \right]-k_{on}\frac{\left[ Ab_{free} \right]}{\varepsilon}\left[ T_{free} \right]+k_{off}\left[ B_{Ab} \right]-k_{e}\left[ T_{free} \right]$$

$$\left( 11 \right) \frac{d\left[ B_{ADC} \right]}{dt}=k_{on}\frac{\left[ ADC \right]}{\varepsilon}\left[ T_{free} \right]-k_{off}\left[ B_{ADC} \right]-k_{e}\left[ B_{ADC} \right]$$

$$\left( 12 \right) \frac{d\left[ B_{ADC,lys} \right]}{dt}=k_{e}\left[ B_{ADC} \right]-k_{deg}[B_{ADC,lys}]$$

$$\left( 13 \right) \frac{d\left[ P_{lys} \right]}{dt}=k_{deg}\left[ B_{ADC,lys} \right]DAR-k_{in}\left[ P_{lys} \right]$$

$$\left( 14 \right) \frac{d\left[ P_{int} \right]}{dt}=k_{in}\left[ P_{lys} \right]+k_{in,p}\left( \frac{1-\varepsilon_{p}}{\varepsilon_{p}} \right)\left[ P_{ext} \right]-k_{out,p}\left[ P_{int} \right]-\frac{k_{on,p}}{\left( 1-\varepsilon_{p} \right)\left( 1+R \right)}\left( P_{target}-\left[ P_{b} \right] \right)\left[ P_{int} \right]+k_{off,p}\left[ P_{b} \right]$$

$$\left( 15 \right) \frac{d\left[ P_{b} \right]}{dt}=\frac{k_{on,p}}{\left( 1-\varepsilon_{p} \right)\left( 1+R \right)}\left( P_{target}-\left[ P_{b} \right] \right)\left[ P_{int} \right]-k_{off,p}\left[ P_{b} \right]$$

$$\left( 16 \right) \frac{d\left[ P_{ext} \right]}{dt}=-k_{in,p}\left( \frac{1-\varepsilon_{p}}{\varepsilon_{p}} \right)\left[ P_{ext} \right]+k_{out,p}\left[ P_{int} \right]$$

$$\left( 17 \right) \frac{d\left[ Ab_{free} \right]}{dt}=-k_{on}\frac{\left[ Ab_{free} \right]}{\varepsilon}\left[ T_{free} \right]+k_{off}[B_{Ab}]$$

$$\left( 18 \right) \frac{d\left[ B_{Ab} \right]}{dt}=k_{on}\frac{\left[ Ab_{free} \right]}{\varepsilon}\left[ T_{free} \right]-k_{off}\left[ B_{Ab} \right]-k_{e}[B_{Ab}]$$

**Tumor Microenvironment Release Drug Dynamics (calculated as a separate cleavage step)**

$$\left( 19 \right) \frac{d\left[ ADC \right]}{dt}=-k_{Cleave}\left[ ADC \right]$$

$$\left( 20 \right) \frac{d\left[ B_{ADC} \right]}{dt}=-k_{Cleave}\left[ B_{ADC} \right]$$

$$\left( 21 \right) \frac{d\left[ Ab_{free} \right]}{dt}=k_{Cleave}\left[ ADC \right]$$

$$\left( 22 \right) \frac{d\left[ B_{Ab} \right]}{dt}=k_{Cleave}\left[ B_{ADC} \right]$$

$$\left( 23 \right) \frac{d\left[ P_{ext} \right]}{dt}=DAR\left( k_{Cleave}\left[ ADC \right]+k_{Cleave}\left[ B_{ADC} \right] \right)$$

**Cell Death Probability**

$$\left( 24 \right) P_{kill}=\frac{P_{\max}\left[ Payload \right]}{Km+[Payload]}$$

$P_{kill}=0 if \left[ Payload \right]<C_{min}$

**Table S1. Baseline PKPD parameters for HER2 and Fc binding**

| **Symbol** | **Parameter** | **Value** | **Unit** | **Source** |
| --- | --- | --- | --- | --- |
| k_10_ | Total antibody clearance | 4.55e-6 | s^-1^ | Singh et al. 2019 [3] |
| k_12_ | Antibody clearance for compartment 1 to 2 | 8.06e-6 | s^-1^ | Singh et al. 2019 |
| k_21_ | Antibody clearance for compartment 2 to 1 | 1.33e-5 | s^-1^ | Singh et al. 2019 |
| P_max_ | Maximum probability of cell killing | 0.045 | - | Calibrated |
| K_m_ | Michaelis-Menten constant | 10 | nM | Calibrated |
| C_min_ | Minimum concentration necessary for cell death | 0.001 | nM | Calibrated |
| P_Target_ | Payload target concentration | 300 | nM | Scheuher et al. 2023 [4] |
| DAR | Enhertu drug-to-antibody ratio | 7.8 | - | Ogitani et al. 2016 [5] |
| k_on_ | ADC binding rate constant | 7.1e-4 | nM^-1^s^-1^ | Bostrom et al. 2011 [6] |
| k_off_ | ADC dissociation rate constant | 3.5e-4 | s^-1^ | Bostrom et al. 2011 |
| K_d HER2_ | HER2 binding affinity | 0.5 | nM | Bostrom et al. 2011 |
| k_e HER2_ | HER2 free internalization rate constant | 3.3e-5 | s^-1^ | Thurber et al. 2007 [7] |
| k_eBound HER2_ | HER2 bound internalization rate constant | 3.3e-5 | s^-1^ | Thurber et al. 2007 |
| K_d Fc_ | Fc binding affinity | 5 | nM | Bostrom et al. 2011 |
| k_e Fc_ | Fc free internalization rate constant | 2.75e-5 | s^-1^ | Mellman et al. 1984 [8] |
| k_eBound Fc_ | Fc bound internalization rate constant | 1.93e-4 | s^-1^ | Mellman et al. 1984 |
| k_deg_ | ADC lysosomal degradation rate constant | 8e-6 | s^-1^ | Maas et al. 2016 [9] |
| ε | Void fraction | 0.24 | - | Schmidt et al. 2009 [10] |
| ε_p_ | Void fraction of payload | 0.44 | - | Bhatnagar et al. 2014 [11] |
| R | Target synthesis rate | 2.75e-11 | M s^-1^ | Thurber et al. 2011 [12] |

**Table S2. Payload PK parameters for T-DXd**

| **Symbol** | **Parameter** | **Value** | **Unit** | **Source** |
| --- | --- | --- | --- | --- |
| k_inPCyt_ | DXd internalization rate from endosome to cytoplasm | 3.16e-3 | s^-1^ | Khera et al. 2018 [13] |
| k_inP_ | DXd cell internalization rate | 3.16e-3 | s^-1^ | Khera et al. 2018 |
| k_outP_ | DXd rate of payload leaving the cell | 2.2e-3 | s^-1^ | Khera et al. 2018 |
| k_onP_ | DXd binding rate to intracellular target | 0.303e-6 | M^-1^s^-1^ | Khera et al. 2018 |
| k_offP_ | DXd unbinding rate to intracellular target | 0.00431 | s^-1^ | Khera et al. 2018 |

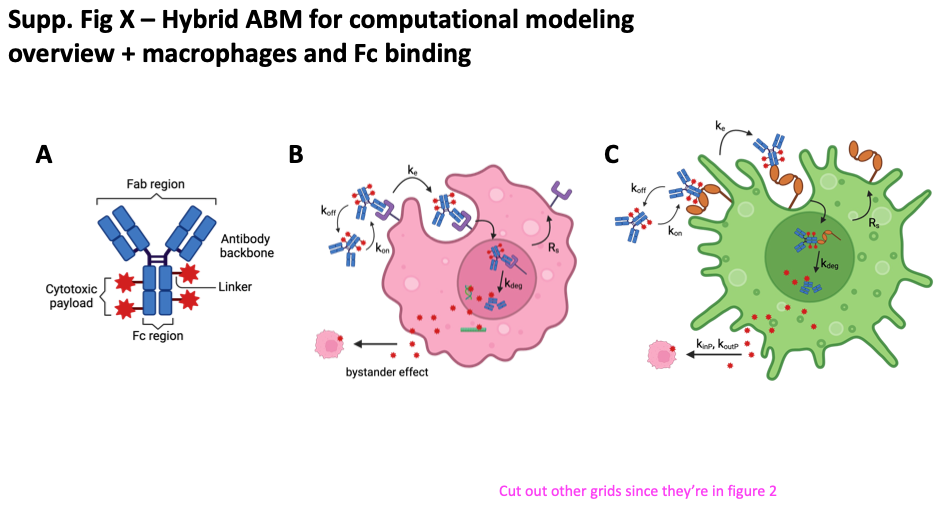

**Figure S1. ADC structure and binding mechanism.**

(A) ADC structure. (B) Primary mechanism of action for ADCs. ADCs bind to the surface of cancer cells via a specific target, and are then internalized and degraded in the cell lysosome to release the payload and cause cell death. (C) ADCs bind to the Fc receptors on macrophages and are internalized to release free payload in the extracellular space. Images generated in BioRender.

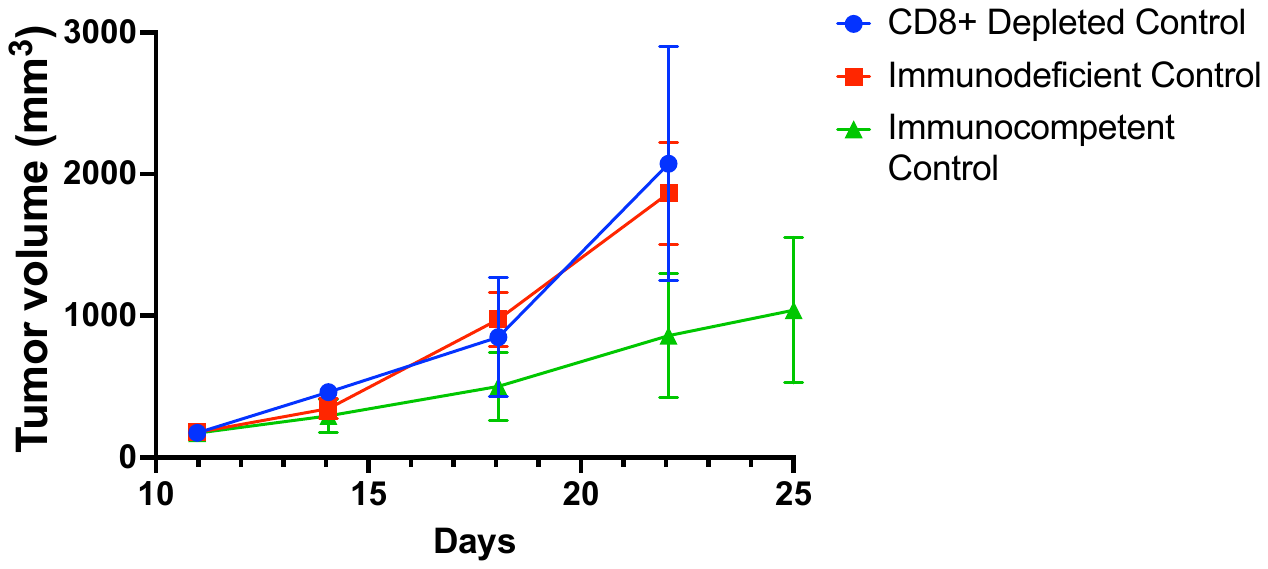

**Figure S2. CT26 control comparison**

We replotted three control tumor growth curves for CT26 cells grown in nude, syngeneic, and CD8+ T cell depleted mouse models from Rios Doria et al. [14] to overlay them for direct comparison. We find that tumor growth is slower in the immunocompetent syngeneic mouse model compared to the immunodeficient nude mouse. However, depleting the CD8+ T cells restores the growth rate to the faster immunodeficient control indicating that CD8+ T cells are the main drivers for slower tumor growth in the syngeneic model.

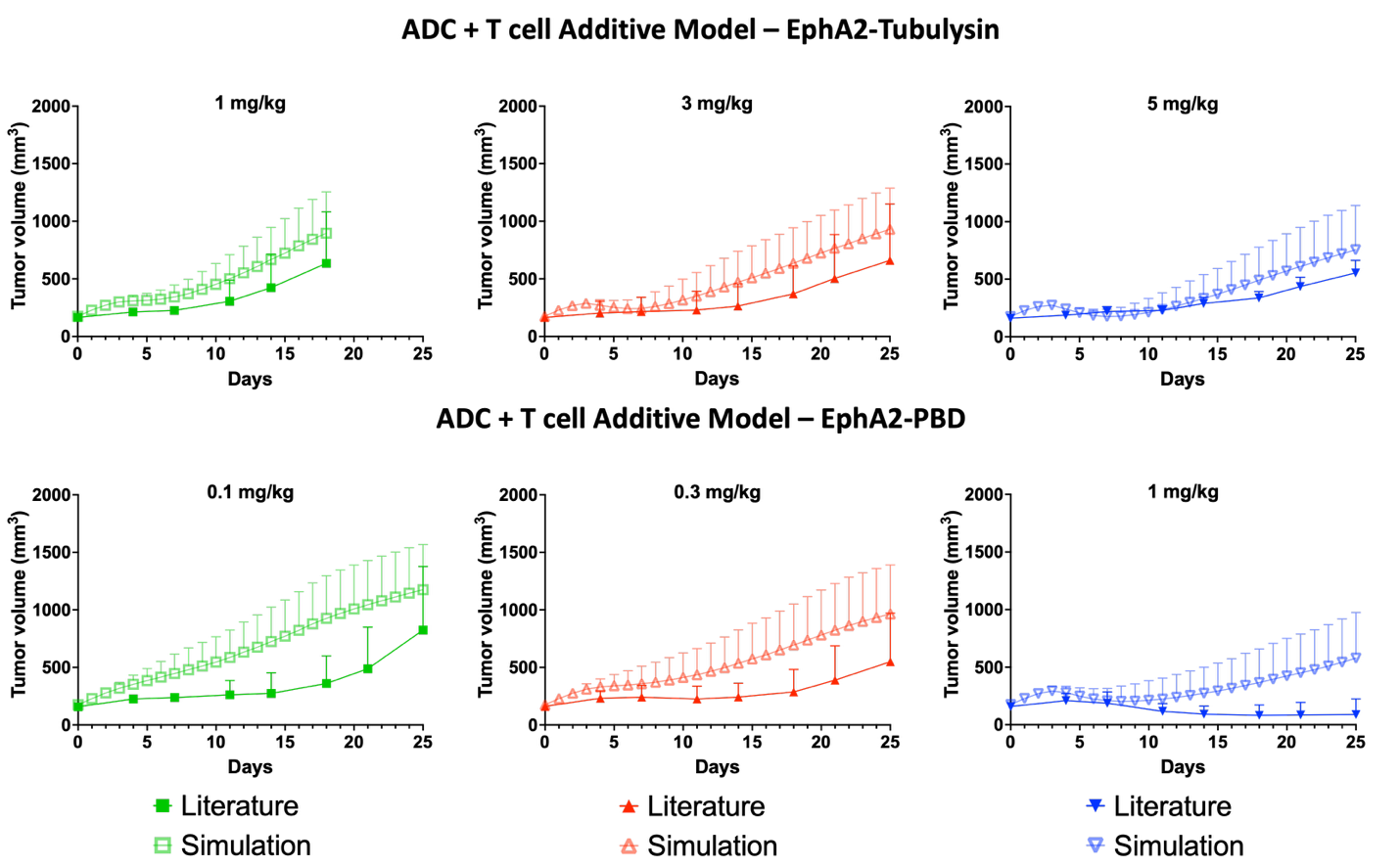

**Figure S3. Additive model simulation results for EphA2-Tubulysin and EphA2-PBD**

After calibrating our PD model parameters to efficacy data for both ADCs treating CT26 nude mice and fitting our T cell killing parameter, we simulated each dose with the baseline level of T cell killing (non-activated) to determine if activation is necessary for efficacy. We find that our additive model under-predicts the efficacy of the syngeneic efficacy data, indicating the necessity for T cell activation for the simulations to match the experimental data. Error bars represent standard deviation; each simulation point is the result of 300 simulation runs (100 simulations run in triplicate).

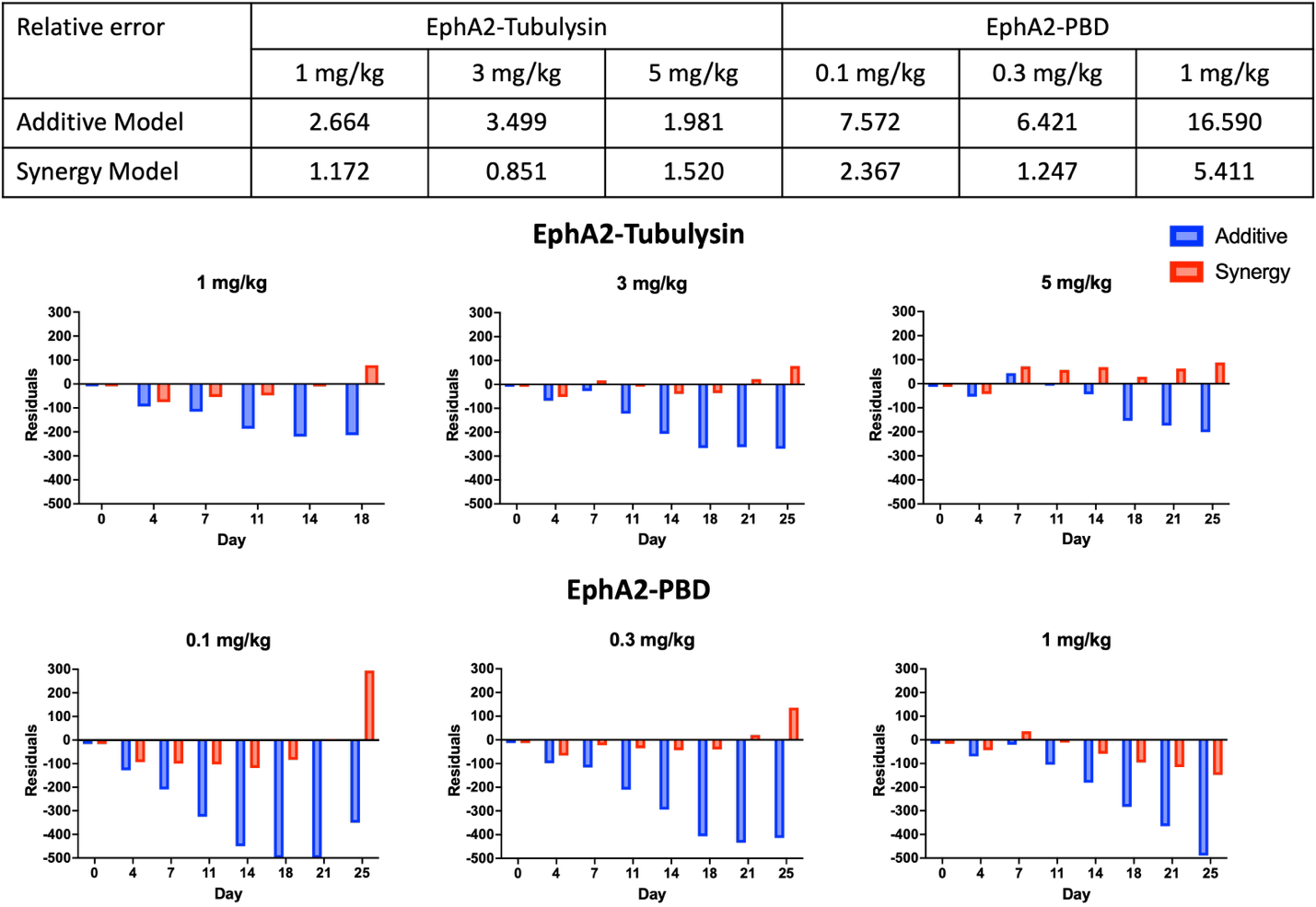

**Figure S4. Comparing relative error and residuals between additive and synergy models**

When we compare the relative errors between the additive and synergy models across all doses of both ADCs, we find smaller errors overall for the synergy model. We also see smaller residuals when plotted over time, visualizing the improved fits of the synergy model fits to the literature data.

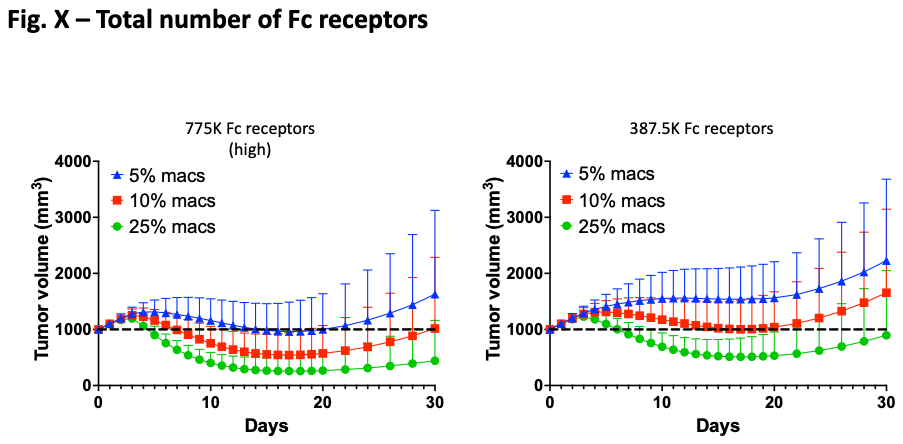

**Figure S5. Total Fc expression impacts T-DXd efficacy**

We varied the percentage of macrophages on the grid with high Fc expression and half Fc expression to determine the impact of varying percentage and Fc expression. With high Fc expression, we still see good efficacy even with 5% macrophages on the grid. Additionally, we see similar efficacies with the same total number of Fc receptors (i.e. 5% with high Fc and 10% with half Fc), indicating that the total number of Fc receptors in the tumor is what drives efficacy. Error bars represent standard deviation; each simulation point is the result of 300 simulation runs (100 simulations run in triplicate).

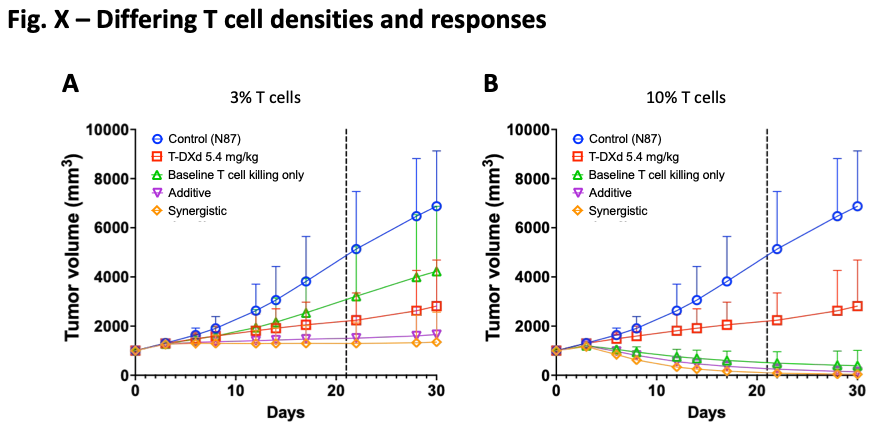

**Figure S6. Enhertu predictions with variable fraction of T cells**

The response rates for the additive (baseline T cell killing plus targeted delivery) and synergy (activated T cell killing plus targeted delivery) models is dependent on the fraction of T cells in the tumor. If we have a smaller fraction of T cells, we see slowed growth with the syngeneic control, but not enough for regression, even with treatment. If we have a larger fraction of T cells we see regression even without treatment. These results highlight the impact of T cells in the tumor, and why many immune-oncology therapies in development focus on immune system activation, particularly with CD8+ T cells. Error bars represent standard deviation; each simulation point is the result of 300 simulation runs (100 simulations run in triplicate).

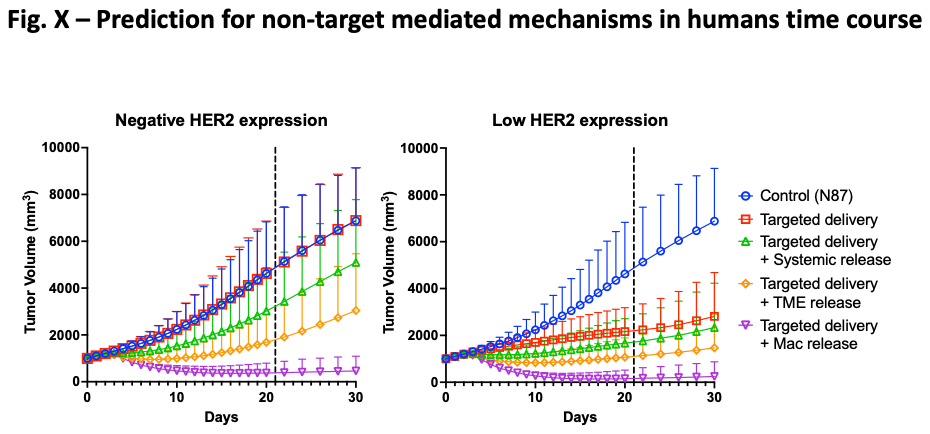

**Figure S7. Time course for each target-mediated and -independent T-DXd mechanism in human tumors**

For comparison, we plotted the tumor volume of each mechanism over 30 days of simulation for our HER2-negative and -low expressing tumors using N87 tumor model growth kinetics. We find that for each mechanism, we see better efficacy for the low expression versus the negative expression, primarily due to the availability of direct targeting sites. Overall, macrophage uptake and payload release contributed the most to tumor regression. Error bars represent standard deviation; each simulation point is the result of 300 simulation runs (100 simulations run in triplicate).

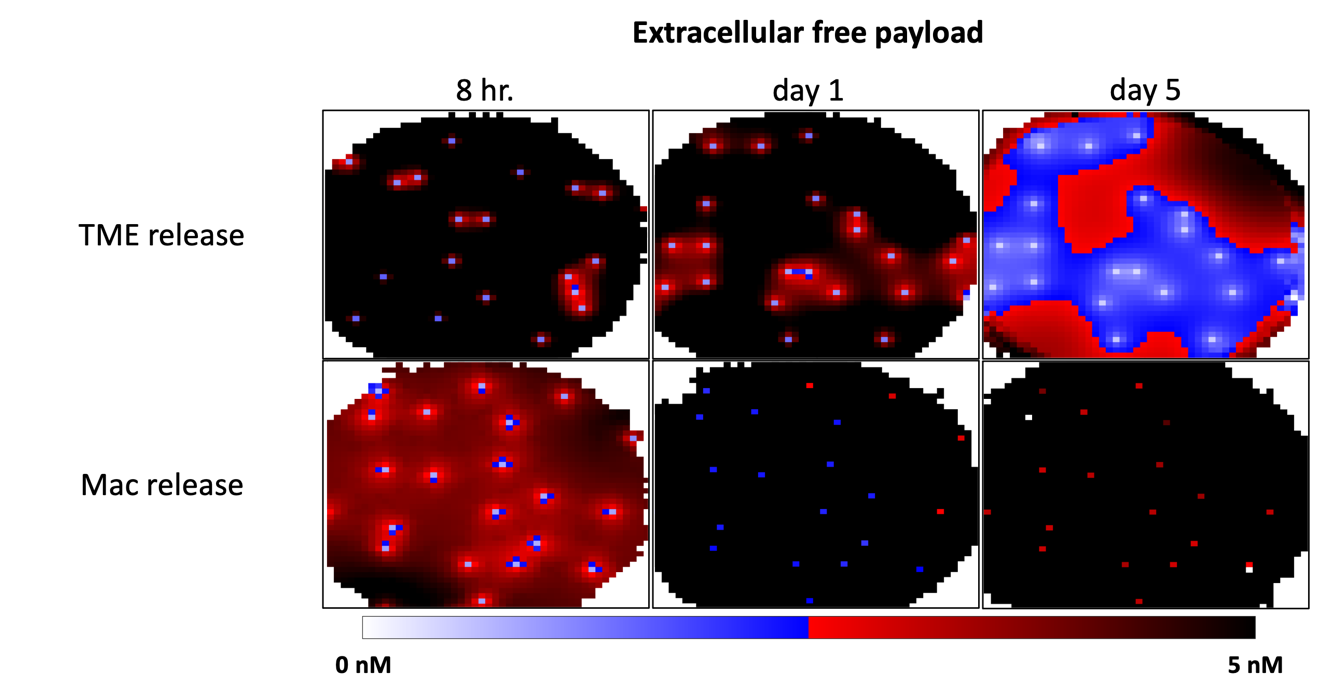

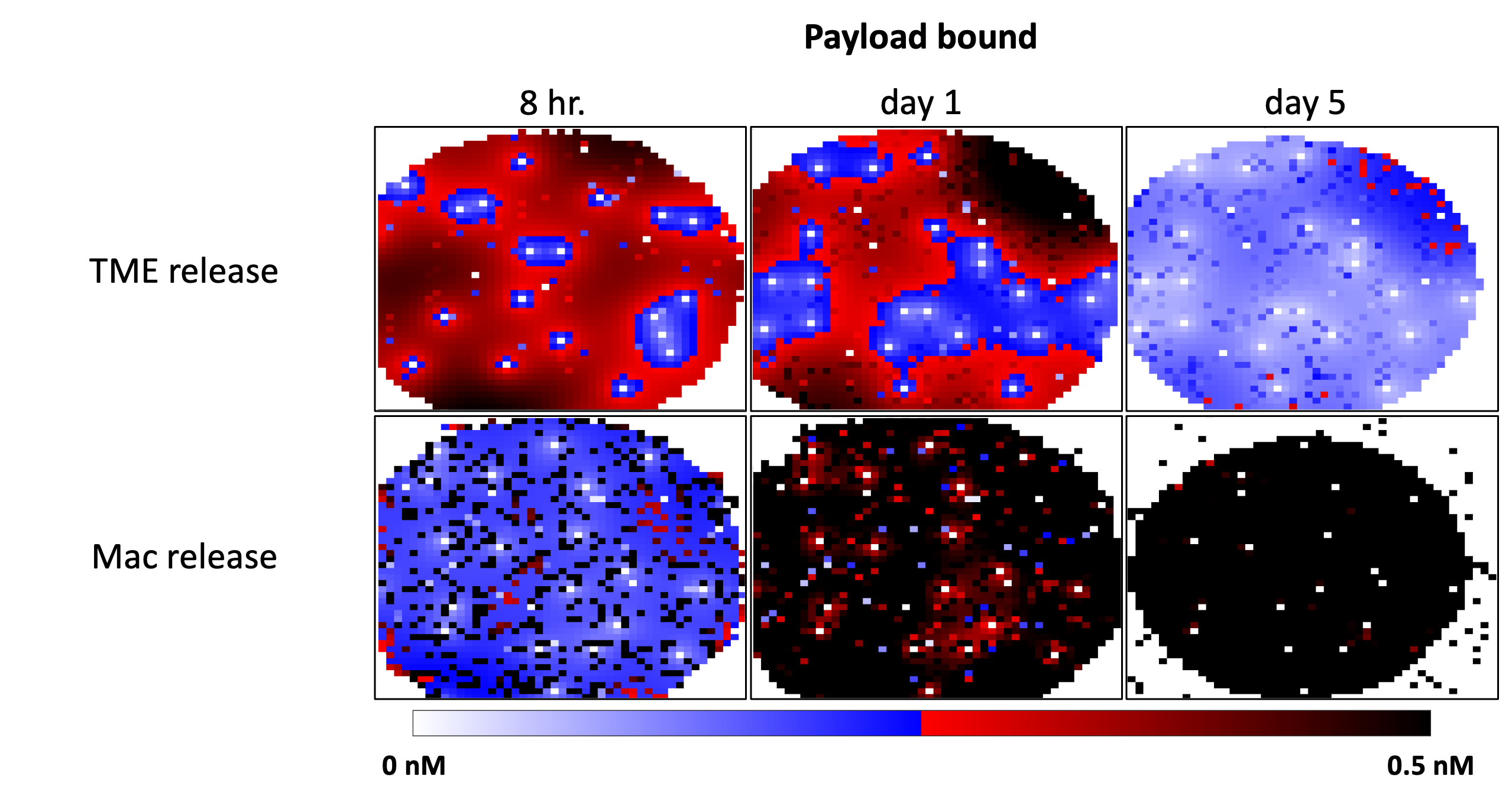

**Figure S8. Distribution time course for extracellular free payload and payload bound between macrophage uptake and extracellular protease cleavage**

At early times (8 hrs), we see increased concentrations for extracellular protease cleavage due to the rapid cleavage rates upon dosing; however free payload is quickly cleared such that by day 5 we no longer have uptake. For macrophages, we see lower concentrations at earlier times due to the trafficking of payload between macrophages and cancer cells, but overall have more sustained free payload in the tumor, even up to day 5. These plots highlight the macrophages ability to retain free payload, leading to better tumor regression.

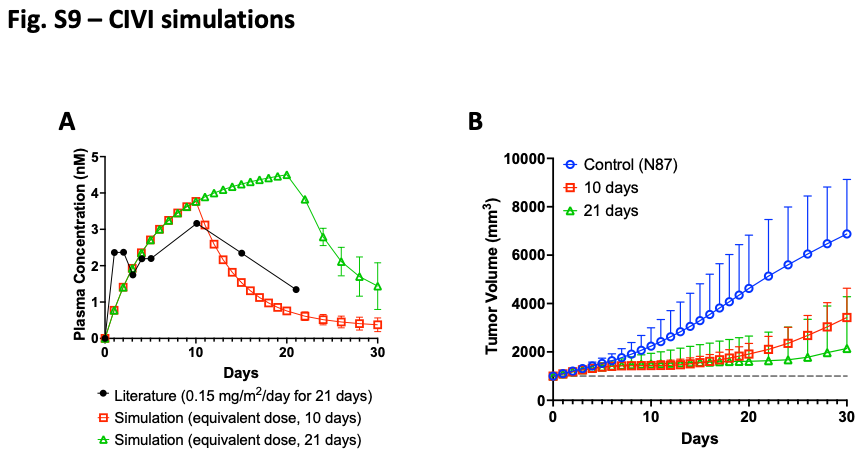

**Figure S9. Free payload PK and efficacy prediction for simulated Continuous Intravenous Infusion (CIVI)**

(A) To evaluate the impact of CIVI on human tumors (with N87 growth kinetics), we matched the PK reported in Garrison et al. [15] for a dose of free exatecan (0.15 mg/m^2^) over 21 days using our previously-used PK parameter set. Though not a perfect fit, we assume that if the simulated concentration of payload in the plasma is higher than the clinically measured values, we’ll be able to see apparent efficacy from free payload. We also simulated the plasma concentration of the same dose over 10 days. (B) For our efficacy curve predictions, we doubled the maximum probability of cell killing to match the potency of exatecan (vs. DXd). When we look at the respective efficacy curve predictions, we do not see tumor regression. Our results highlight that it is less likely that efficacy from dosing Enhertu in HER2-low and -negative expression systems comes from free payload in the plasma since even sustained levels at a similar concentrations of a more potent payload did not achieve significant efficacy in the clinic. Error bars represent standard deviation; each simulation point is the result of 300 simulation runs (100 simulations run in triplicate).

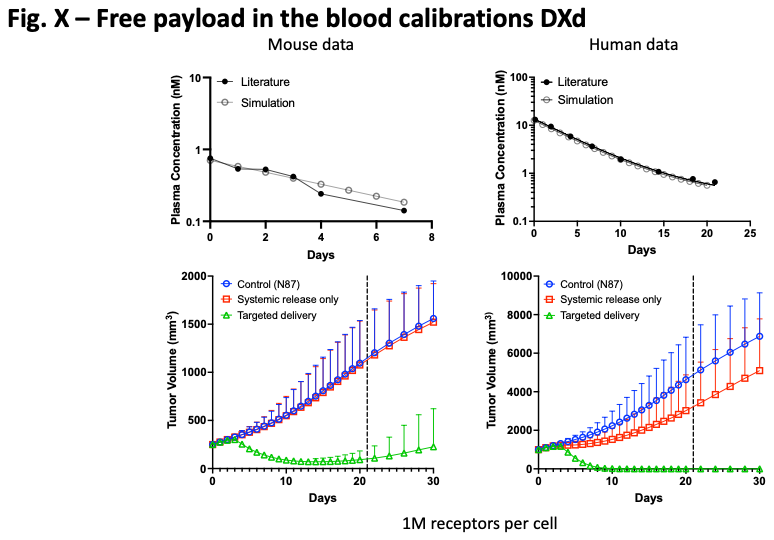

**Figure S10. Mouse and human free payload PK calibration and efficacy prediction for T-DXd**

To evaluate the impact of free payload in the blood after dosing T-DXd, we fit the PK for mouse [16] and human [17] data to get the clearance rates and equivalent dose of free payload to match the initial concentrations. We simulated this dose of free payload in N87 tumors (1M receptors/cell) to see their impact against the clinical dose of T-DXd (5.4 mg/kg), and found minimal impact from the equivalent free payload dose. Error bars represent standard deviation; each simulation point is the result of 300 simulation runs (100 simulations run in triplicate).

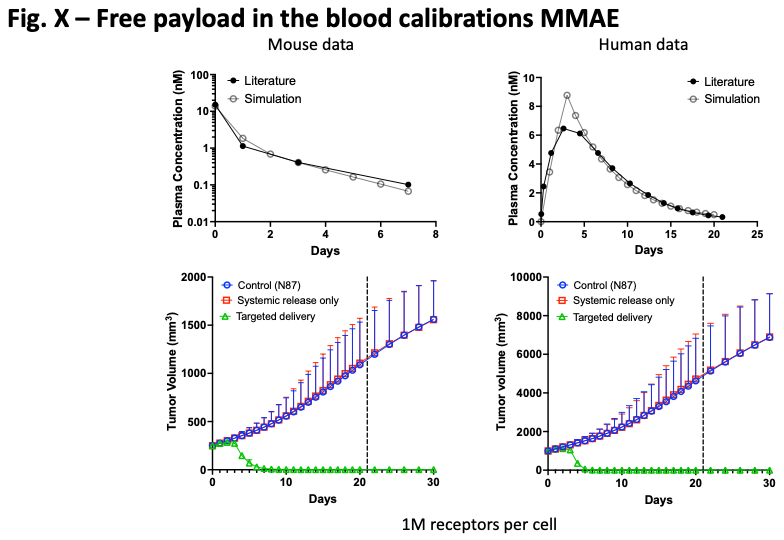

**Figure S11. Mouse and human free payload PK calibration and efficacy prediction for T-MMAE**

To evaluate the impact of free payload in the blood after dosing T-MMAE, we fit the PK for mouse [3] and human [17] data to get the clearance rates and equivalent dose of free payload to match the initial concentrations. We simulated this dose of free payload in N87 tumors (1M receptors/cell) to see their impact against the reported dose of T-MMAE (10 mg/kg), and found essentially no impact from the equivalent free payload dose. Error bars represent standard deviation; each simulation point is the result of 300 simulation runs (100 simulations run in triplicate).

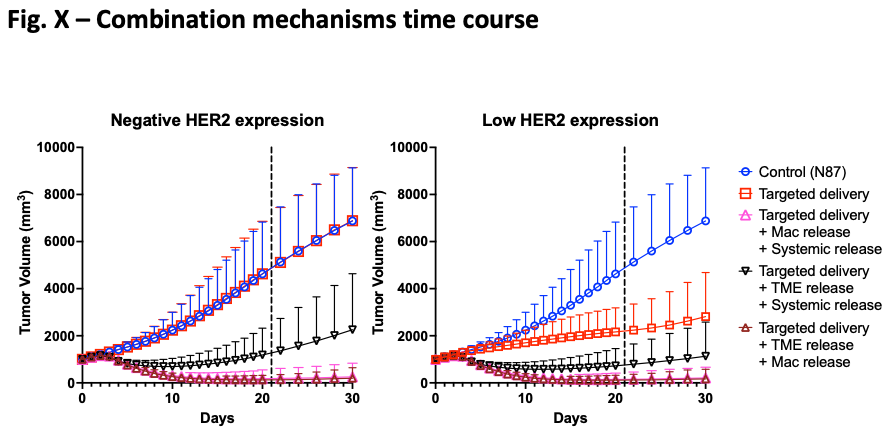

**Figure S12. Time course for combined target-mediated and -independent T-DXd mechanisms in N87 tumors**

We simulated a series of combined mechanisms to determine the impact multiple mechanisms of action on T-DXd efficacy. The addition of free plasma payload to extracellular cleavage is enough for tumor regression, and we see further regression when macrophage uptake is combined with either free plasma payload or extracellular cleavage. Error bars represent standard deviation; each simulation point is the result of 300 simulation runs (100 simulations run in triplicate).

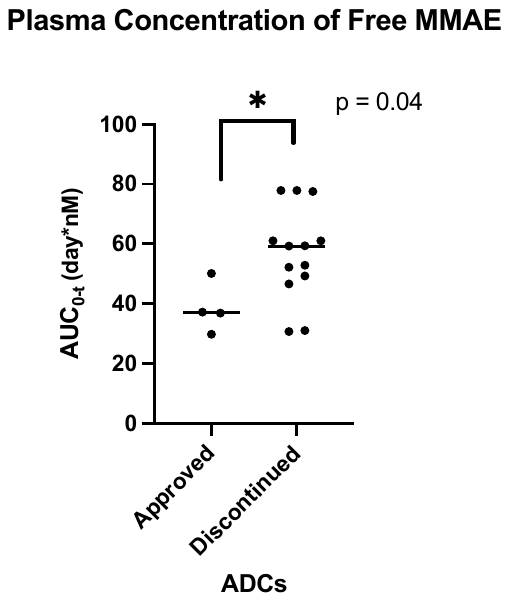

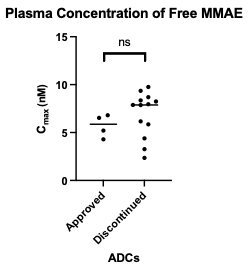

**Figure S13. Plasma concentration comparisons between approved and discontinued ADCs with MMAE payloads**

When comparing the plasma concentrations for MMAE-based ADCs as outlined in Chang et al. [18], for C_max_, we find no significant difference between approved and discontinued ADCs. For AUC, we see a statistically significant lower concentration of MMAE for approved drugs, indicating lower payload exposure from the plasma for approved ADCs.
